# Characterization of Tgl2, a putative lipase in yeast mitochondria

**DOI:** 10.1101/2024.05.08.593122

**Authors:** Vitasta Tiku, Takashi Tatsuta, Martin Jung, Doron Rapaport, Kai Stefan Dimmer

## Abstract

Mitochondria derive the majority of their lipids from other organelles through contact sites. These lipids, primarily phosphoglycerolipids, are the main components of mitochondrial membranes. In the cell, neutral lipids like triacylglycerides (TAGs) are stored in lipid droplets, playing an important role in maintaining cellular health. Enzymes like lipases mobilize these TAGs according to cellular needs. Neutral lipids have not yet been reported to play an important role in mitochondria so the presence of a putative TAG lipase – Tgl2, in yeast mitochondria is surprising. Moreover, *TGL2* and *MCP2*, a high-copy suppressor for ERMES deficient cells, display genetic interactions suggesting a potential link to lipid metabolism. In this study, we characterize in detail Tgl2. We show that Tgl2 forms dimers through intermolecular disulfide bridges and a cysteine-dependent high molecular weight complex. Furthermore, we could identify the lipase motif and catalytic triad of Tgl2 through *in silico* comparison with other lipases and mutated the catalytically active residues accordingly. Both mutants failed to rescue the growth phenotype of *mcp2*Δ*/tgl2*Δ double deletion strain suggesting that the residues are indeed essential for the protein’s function. Additionally, we discovered that the catalytically active aspartate residue is important for protein stability. Steady state level analyses with non-functional variants of Tgl2 led to the identification of Yme1 as the protease responsible for its quality control. Finally, we provide evidence that the overall increase in TAGs in cells lacking Mcp2 and Tgl2 originates from the mitochondria. Collectively, our study provides new insights into a key player in mitochondrial lipid homeostasis.

## Introduction

Mitochondria are complex organelles with a wide array of functions. Their endosymbiotic bacterial ancestry has enabled them to evolve into semi-autonomous, bilayered entities consisting of an outer (MOM) and an inner mitochondrial membrane (MIM) (1, 2). These membranes are distinct in their protein content and lipid composition, separating the organelle into two aqueous sub-compartments – the matrix and the intermembrane space (IMS) (1).

Mitochondrial membranes have a dynamic architecture which is enabled by a continuous supply of lipids and proteins. While some lipids like cardiolipin are synthesized *in organello* from its precursor phosphatidic acid, mitochondria predominantly acquire lipids from the ER. Contact sites facilitating lipid exchange between mitochondria and lipid droplets (LDs), have also been suggested (3, 4). Once imported, these lipids need to be re-distributed between the two membranes, but the exact mechanism for this is still unknown. Alterations in the MICOS (mitochondrial contact site) complex have been shown to disturb mitochondrial ultrastructure and disrupt cristae morphology, suggesting a potential contribution to lipid metabolism (5–7). Further, lipid transfer proteins (LTPs) mediating a constant exchange of precursor lipids between the MIM and MOM, by shuttling them across the IMS have also been reported (8, 9).

The IMS is a small and rather crowded sub-compartment, housing ~50 proteins in yeast and humans (10, 11). While these numbers account for the soluble protein population, there is also a considerable fraction of MIM and MOM proteins that protrude into the IMS. As a buffer between the two membranes, the IMS allows for exchange of lipids, metabolites, and movement of proteins to maintain mitochondrial function. These proteins are typically small and are imported through either the disulfide relay pathway or the stop-transfer pathway. Additionally, some proteins such as CytC, Prx1, and Adk1 to name a few, are imported through atypical pathways some of which are yet to be fully elucidated (12, 13).

Tgl2 is a TAG lipase localized to the mitochondrial intermembrane space (14, 15). It belongs to the Tgl family of lipases in *S. cerevisiae* of which Tgl1, Tgl3, Tgl4, and Tgl5 can be found in LDs or cytosol (16). A previous study suggested lipolytic activity of Tgl2 towards short-chain TAGs and DAGs and long-chain TAGs (14). The protein has also been shown to compensate for the absence of DAG Kinase in *E. coli* (17). While deletion of *TGL2* does not result in a detectable phenotype, we previously reported negative genetic interactions of *TGL2* with *MCP2* (15). Elevated levels of Mcp2 appear to maintain lipid trafficking to and from mitochondria in ERMES (ER mitochondria encounter structure) deficient yeast (18), and the protein was more recently suggested to be involved in CoQ mobilization from the MIM (19).

Phosphatidylethanolamine (PE) and cardiolipin (CL), which is exclusive to mitochondria, are synthesized in mitochondria from precursor phosphoglycerolipids (PGL) that not only need to be imported but also have to be transported across the IMS, since certain biosynthesis steps take place in the MIM. PE is synthesized from phosphatidylserine (PS) by the MIM decarboxylase Psd1 (20). CL is assembled from the precursor phosphatidic acid (PA) in the MIM, before maturation of the unique lipid takes place in the MOM (9). Of the repertoire of IMS proteins, the only evolutionarily conserved lipid transfer system is the Ups1/Ups2/Mdm35 complex (9, 21–23). This system is involved in the trafficking of the aforementioned lipids PA and PS (9, 20, 22, 23). Therefore, and since neutral lipids like TAGs are primarily stored in and mobilized from LDs, reports on the presence of a putative TAG lipase in the IMS are of high interest.

In this study, we characterized Tgl2 and study its quality control. We identified its putative catalytic site and investigated its effect on cellular neutral lipid content.

## Results

### Tgl2 forms a homodimer that is sensitive to reducing agents

Tgl2 harbours 8 cysteine residues that are essential for its import through the Mia40 pathway (15). Since all our previous studies about Tgl2 were performed under reducing conditions, we asked whether the protein would behave differently under non-reducing conditions, especially since the IMS has a more oxidizing environment than the cytosol or matrix (24). We isolated crude mitochondria from yeast cells over-expressing N-terminally HA tagged Tgl2 (HA-Tgl2) and omitted any reducing agents during SDS-PAGE. A higher molecular weight species, approximately twice the size of Tgl2, was observed (Fig. 1A). Further, this species was lost over time as it could be detected more evidently in whole cell lysates and crude mitochondria isolated after mechanical rupturing but to a lesser extent in mitochondria obtained from spheroplasting-based isolation protocols (Fig. 1B). We further quantified this higher molecular weight species using the MOM protein Tom20 as a loading control. We observed that after shorter isolation protocols, this species contributes to almost half of the total protein whereas its appearance is noticeably reduced upon longer isolation protocols (Fig. 1B). Therefore, we assume that the species is not a consequence of oxidation, which would be expected to increase over time.

**Fig. 1.**
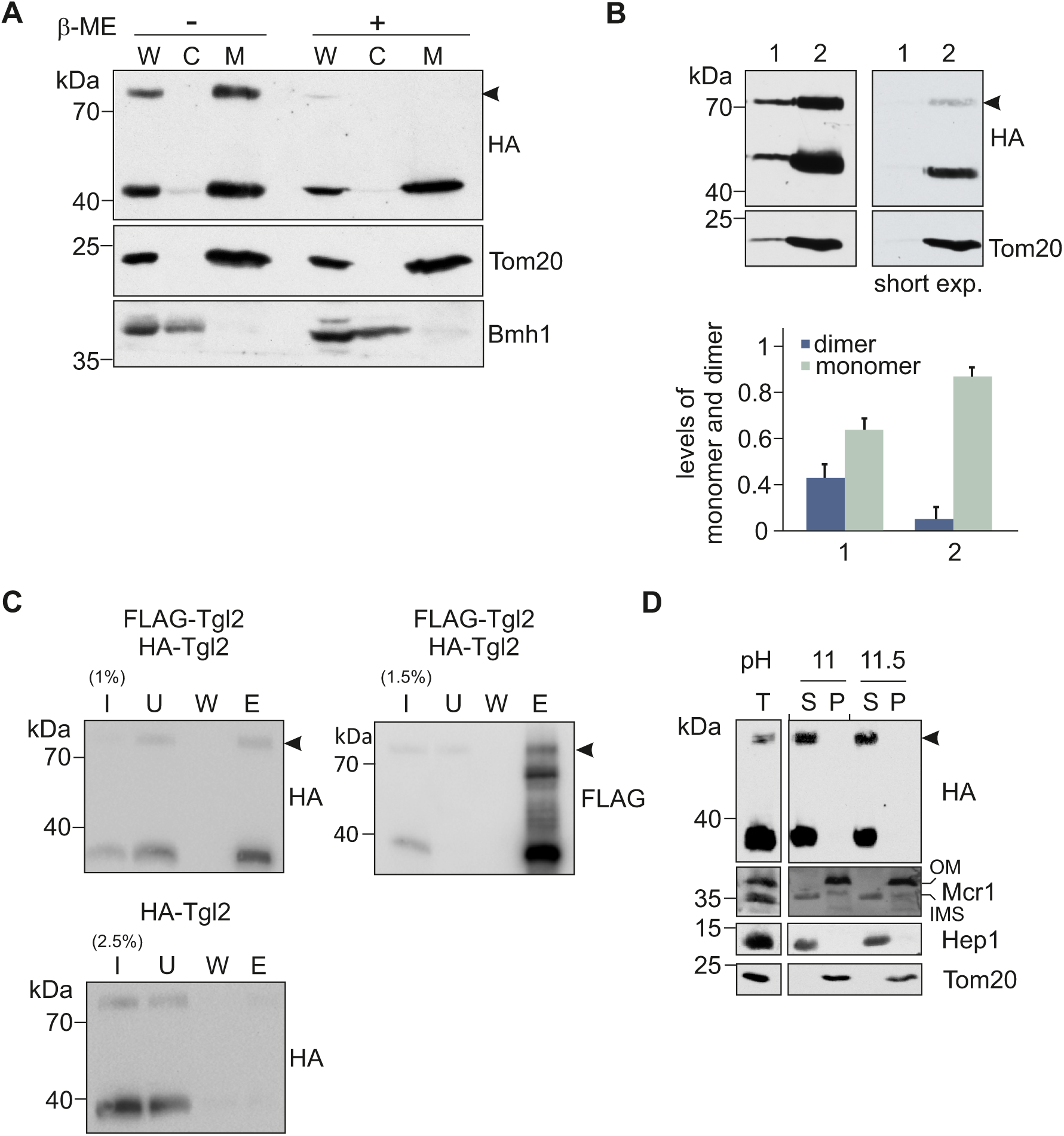
Tgl2 forms a homodimer that is sensitive to reducing agents. (A) Tgl2 forms a higher molecular weight adduct under non-reducing conditions. Cells lacking *TGL2* (*tgl2*Δ) were transformed with a plasmid encoding HA-Tgl2 and grown to mid-logarithmic phase. Mechanical lysis was used to isolate whole cell (W), cytosolic (C), and mitochondrial (M) fractions. The samples were mixed with sample buffer with or without β-mercaptoethanol (β –ME) and analysed by SDS-PAGE and immunodecoration with antibodies against the HA-tag, as well as Tom20 (mitochondrial marker) and BmhI (cytosolic marker). The higher molecular weight adduct is marked by an arrowhead. (B) The higher molecular weight adduct is more pronounced upon shorter isolation protocols. Cells expressing HA-Tgl2 were grown to mid-logarithmic phase and mitochondrial fractions were isolated afterwards either mechanical lysis (1) or spheroplasting (2). The samples were analysed by non-reducing SDS-PAGE and immunodecoration. Quantification of the adduct was done with Tom20 as a loading control and he total Tgl2 amounts were taken as 100%. The graph represents the mean values ± SD of four independent experiments. (C) The adduct is a homodimer of Tgl2. Mitochondria were isolated from *tgl2*Δ cells expressing either HA-Tgl2 or co-expressing HA-Tgl2 and FLAG-Tgl2. The organelles were solubilized and subjected to co-immunoprecipitation with anti-FLAG beads. Samples from input (I), unbound (U, 5%), wash (W, 5%) and eluate fractions (E, 50%) were analyzed by SDS-PAGE and immunodecoration with antibodies against the FLAG-tag. (D) Tgl2 dimer is soluble. Mitochondria containing HA-Tgl2 were subjected to alkaline extraction at different pH values. The supernatant (S) and pellet (P) fractions were analysed by SDS-PAGE and immunodecoration with the indicated antibodies. Tom20 (membrane integrated protein), Hep1 (soluble matrix protein), Mcr1 (a protein with two isoforms, long 34 kDa form embedded in the MOM and shorter soluble form in the IMS).

Since the size of this higher molecular weight species is twice that of Tgl2, we assumed that it could be a homodimer of Tgl2. To test our hypothesis, we isolated mitochondria containing two variants of Tgl2 – N-terminally tagged with either HA- or FLAG-tag and performed co-immunoprecipitation with beads containing antibodies against the FLAG-tag. We analysed the fractions by Western blotting and immunodecoration with antibodies against the two different tags. HA-tagged Tgl2 was detected only in the eluate fraction from mitochondria containing both tagged proteins. As a control, we could verify that the anti-FLAG beads do not unspecifically interact with HA-Tgl2. Furthermore, we could also detect the higher molecular weight species of both variants in the eluate fraction (Fig. 1C, arrowhead). Taken together, the two tagged Tg2 species are indeed interacting with each other, and the higher molecular species is a homodimer.

The submitochondrial localization of Tgl2 was previously determined through hypo-osmotic lysis assays. We previously showed by alkaline extraction that monomeric Tgl2 is not embedded in a mitochondrial membrane (15). However, we wondered if varying the pH could alter the response of the either the monomeric or the dimeric forms of the protein to alkaline extraction, by retaining any membrane interactions. Isolated mitochondria were subjected to alkaline extraction assay and the pellet and supernatant fractions were analysed by SDS-PAGE and immunodecoration was performed with Tom20, Mcr1, and Hep1 as controls for membrane integrated and soluble proteins (Fig. 1D). We observed that the dimeric form is also found in the soluble fraction, indicating that both forms of the protein are soluble.

### Tgl2 is associated with the mitochondrial inner membrane

A protein like Tgl2, with a suggested affinity for lipids, would need to bind to either of the mitochondrial membranes to gain access to potential substrates. We asked whether we could observe a specific interaction of Tgl2 with vesicles of one of the mitochondrial membranes. To ascertain this, we performed a sucrose gradient centrifugation with mitochondrial vesicles containing HA-Tgl2 (Fig. 2A). The fractions of the gradient with increasing density were analysed by Western blotting. Tgl2 was found in vesicles of higher density that also contained the *bona fide* MIM protein Cox2 and was almost completely absent in lighter fractions marked by the presence of the MOM protein Tom20 (Fig. 2B and C). These observations suggest that the vast majority of Tgl2 tends to associate with the MIM.

**Fig. 2.**
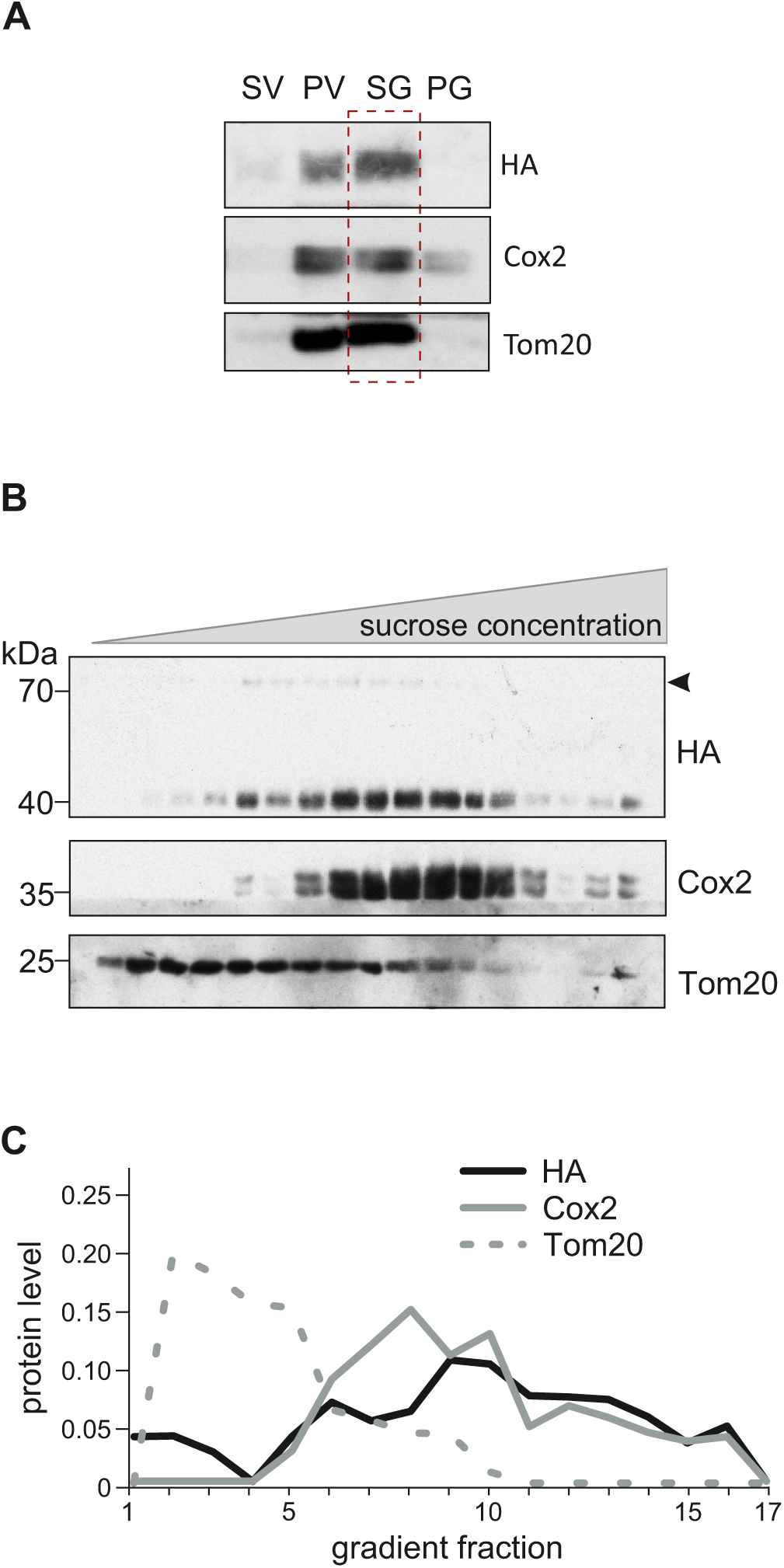
Tgl2 associates with the mitochondrial inner membrane. (A) The majority of Tgl2 is found in mitochondrial vesicles. Mitochondrial vesicles were obtained after swelling and sonication of mitochondria from cells containing HA-Tgl2. The indicated control fractions were collected after swelling, vesicle generation, and clarifying spins. The sample was sonicated to form vesicles followed by clarifying spin to yield supernatant – SV and pellet – PV fractions. The vesicles in PV was sonicated again and subjected to a slower clarifying spin resulting in SG (supernatant) and PG (pellet). Fraction SG was further subjected to a sucrose gradient to separate the MOM and MIM vesicles. The fractions were analysed by SDS-PAGE and immunodecoration. (B) Tgl2 associates with MIM vesicles. The mitochondrial vesicles from S4 fraction (depicted in panel A) were separated by sucrose density centrifugation. Fractions were collected and analysed by SDS-PAGE and immunodecorated with the indicated antibodies, Cox2 (MIM protein) and Tom20 (MOM protein). (C) Tgl2 and Cox2 containing fractions coincide. Intensities corresponding to Tgl2, Cox2, and Tom20 were quantified and normalized to the total content of each protein.

### Tgl2 is a component of a higher molecular weight complex

We previously identified *TGL2* in a synthetic genetic array as a negative interactor of *MCP2* - encoding the kinase Mcp2, an IMS facing protein involved in mitochondrial lipid and coenzyme Q homeostasis (15, 18, 19). Although we tried several different experimental approaches, we could not observe any physical interaction between Tgl2 and Mcp2. (data not shown). Furthermore, co-immunoprecipitation experiments with tagged Tgl2 (HA-Tgl2) and subsequent proteomic analysis failed to reveal Mcp2 as a putative interactor. Of note, no other prominent putative interaction partner(s) could be identified through proteomics analysis (data not shown). The proteomics analysis revealed as hits, ribosomal proteins and abundant mitochondrial proteins such as Porin and Om45. The dataset also contained MIM proteins like Mir1, Cox2 and Cyc1.

Even though we could not identify any promising putative interactors, we wanted to know if Tgl2 forms a higher molecular weight complex. To address this, we isolated mitochondria from yeast overexpressing HA-Tgl2, solubilized them with either digitonin or Triton X-100, and subjected them to blue native (BN)-PAGE followed by immunodecoration. We observed that Tgl2 migrates as part of a complex of 500kDa that was more stable upon mild solubilization using digitonin (Fig. 3A). As expected, and to verify that the anti-HA does not display any cross-reactivity, no signal was detected in cells transformed with an empty vector. This approach was also used to analyse the Tgl2-complex in yeast deletion strains of potential physical interactors selected by educated guess. Table S1 provides an overview of the different deletion mutants analysed and the roles of the respective proteins in mitochondrial physiology. We did not observe a reduction in size or loss of the Tgl2-complex in any of these mutants. We also monitored the steady state levels of monomeric and dimeric Tgl2 in the various deletion strains and the dimeric protein was always detected (Fig. S1). Of note, the Tgl2 complex appears as a sharp solitary band unlike known mitochondrial membrane embedded complexes like TOM complex that displays a rather broad band in native gels (Fig. 3A). The precise apparent complex of Tgl2 is supported also by a recent high throughput complexome study (25). This behaviour further suggests that Tgl2 is indeed a soluble protein of defined quaternary structure without associated membrane lipids.

**Fig. 3.**
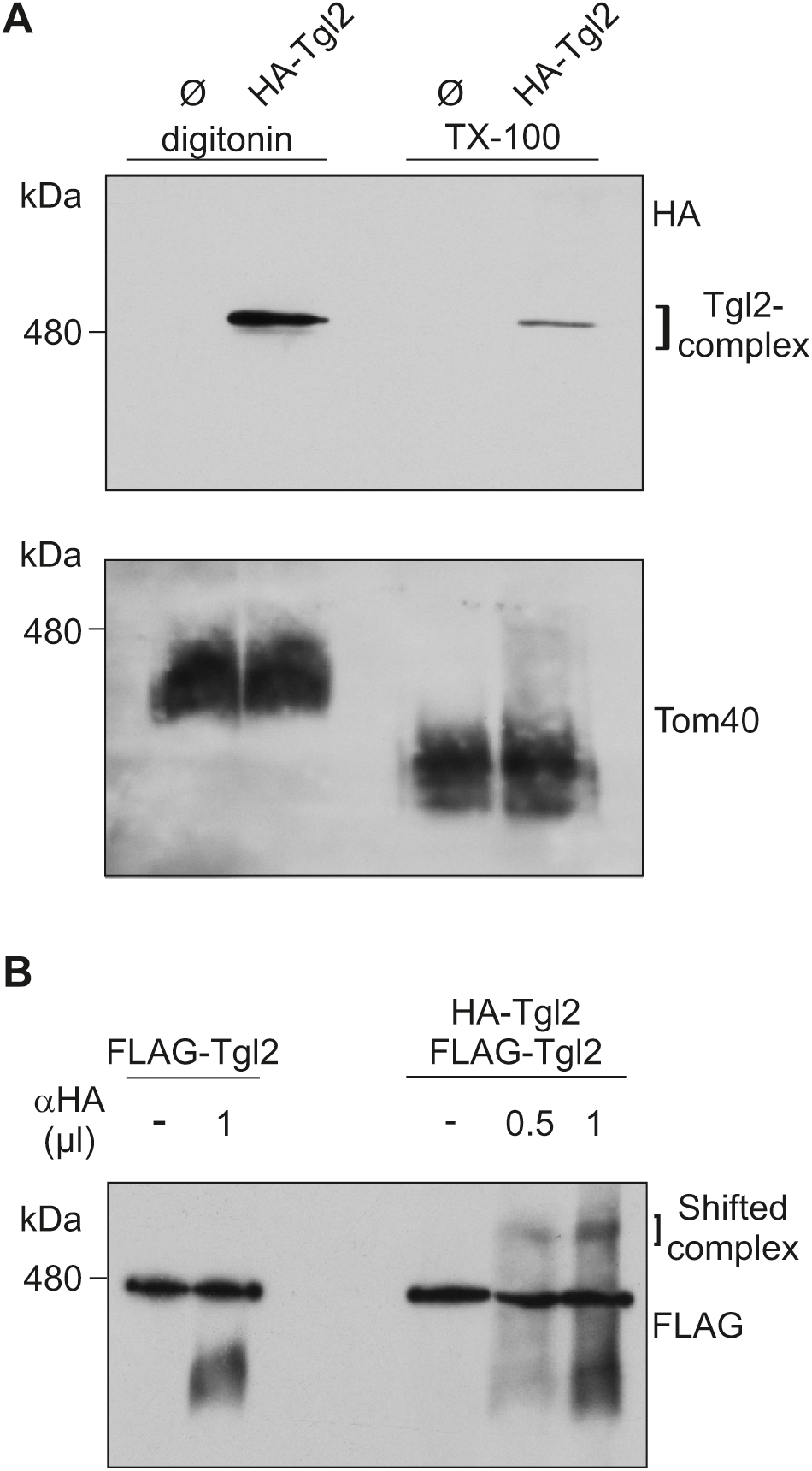
Tgl2 is a component of a higher molecular weight complex. (A) Tgl2 forms a higher molecular weight complex. Mitochondria containing HA-Tgl2 were solubilized with either digitonin or TritonX-100 (Tx-100) and subjected to BN-PAGE (4-14%) and immunodecoration with antibodies against either HA-tag or Tom40, as a loading control. (B) Tgl2 complex contains at least 2 copies of the protein. Mitochondria containing either only HA-Tgl2 or HA-Tgl2 and FLAG-Tgl2 were solubilized with digitonin. The lysate was incubated with 0, 0.5, or 1 µL anti-HA antibody (α-HA) and analysed by BN-PAGE (4-14%) and immunodecoration with anti-FLAG antibody.

We next asked if the Tgl2-containing complex contains multiple copies of the protein. To address this, we performed an antibody shift assay with solubilised mitochondria containing one or two differently tagged variants of Tgl2. Prior to the BN-PAGE analysis we incubated the mitochondrial lysate with an antibody against the HA-tag and immunodecorated against the FLAG-tag. We observed a size shift only for the sample containing both tagged Tgl2 variants (Fig. 3B). In contrast, the complex was not shifted when mitochondria containing only FLAG-Tgl2 were used. These observations indicate that the complex contains at least two copies of Tgl2.

### The MIM protease Yme1 mediates the quality control of Tgl2

Previous studies regarding quality control of IMS proteins have reported that Yme1 – an iAAA protease of the MIM, is responsible for degrading misfolded or unstable proteins in the IMS (26, 27). Many of these proteins are also substrates of the disulfide relay import pathway. We previously observed that C-terminally modified Tgl2 and Tgl2_Cys_, whose cysteine residues were mutated to serine, were hardly detectable in WT yeast cells. These protein variants were also unable to rescue the growth defect exhibited by the *mcp2*Δ/*tgl2*Δ double mutant (15). To test whether non-functional and potentially misfolded Tgl2 variants are substrates of Yme1, we compared the steady state levels of N-terminally tagged Tgl2 (HA-Tgl2), C-terminally tagged Tgl2 (Tgl2-HA), and a variant of Tgl2 wherein all the cysteines are mutated to serine (HA-Tgl2_cys_) in *tgl2*Δ to those in *yme1*Δ cells. We previously reported that only a functional variant of Tgl2 can rescue the growth phenotype exhibited by *mcp2Δ/tgl2*Δ, and consequently used this as a system to assess the functionality of HA-Tgl2, Tgl2-HA, and HA-Tgl2_cys._ We performed a drop-dilution assay on non-fermentable (glycerol) carbon sources and could confirm these observations (Fig. 4A and (15)). A stark growth defect was observed for *mcp2Δ/tgl2*Δ on glycerol-containing medium which could only be rescued by introducing HA-Tgl2 but not the other two variants, further confirming that Tgl2-HA and HA-Tgl2_cys_ are non-functional. To compare the steady state levels of these variants, we obtained whole cell lysate, cytosolic, and crude mitochondrial fractions from either *tgl2*Δ or *yme1*Δ expressing these protein variants, and analysed them Western blotting (Fig. 4B). We could detect the native-like HA-Tgl2 in *tgl2*Δ as well as *yme1*Δ cells. However, Tgl2-HA and HA-Tgl2_cys_ were only detected in the fractions from *yme1*Δ cells. These findings confirm the notion that Yme1 is involved in the elimination of non-functional variants of Tgl2. Interestingly, HA-Tgl2_cys_ is unable to form a homodimer, strengthening the hypothesis that the observed dimer is indeed due to disulfide bridges.

**Fig. 4.**
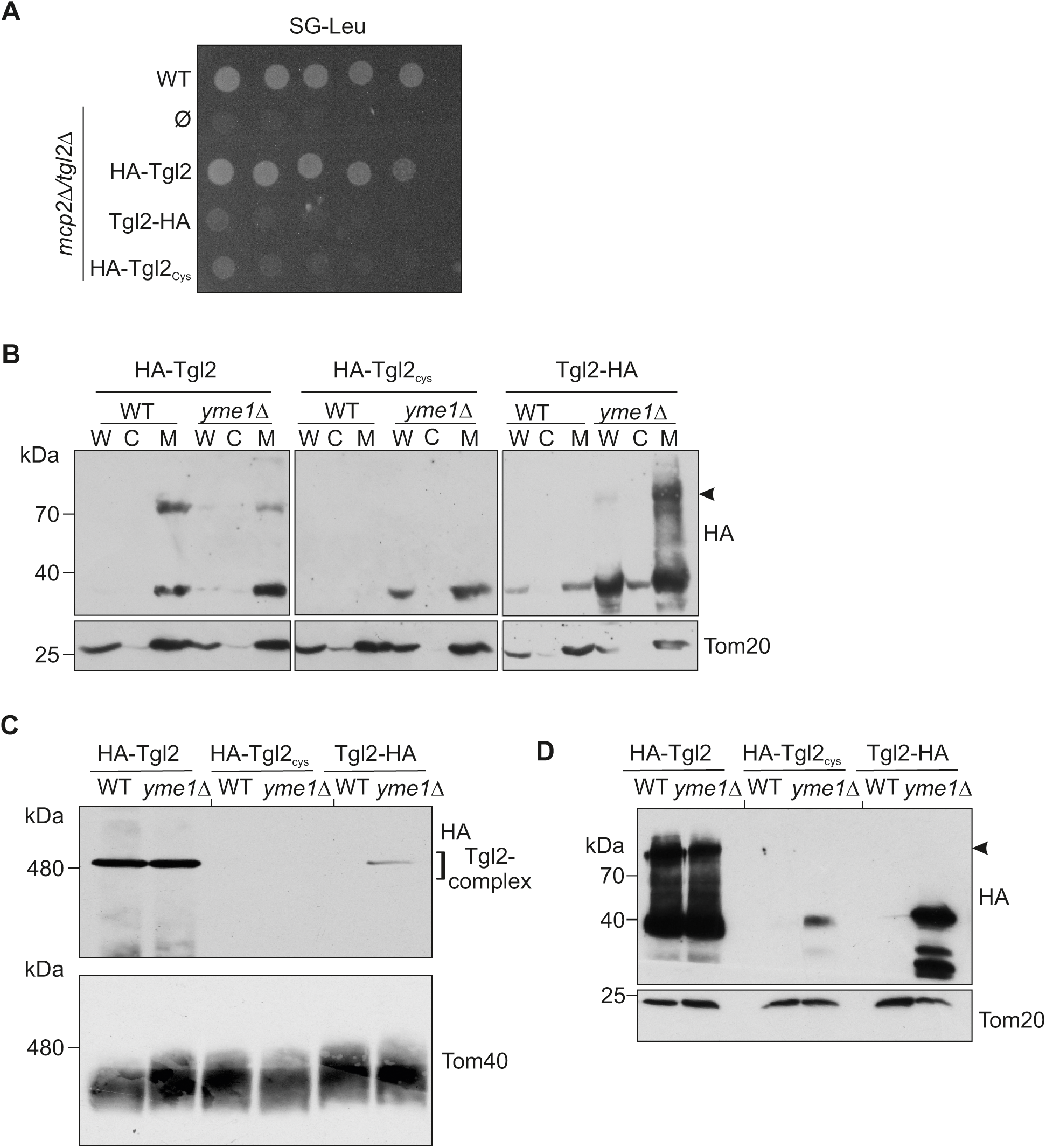
The MIM protease Yme1 mediates the quality control of Tgl2. (A) C-terminally tagged Tgl2 and Tgl2 variant with all its cysteines mutated to serine do not rescue the growth defect of *mcp2*Δ*/tgl2*Δ. WT or *mcp2*Δ*/tgl2*Δ cells were transformed with an empty vector (Ø) or a plasmid encoding the indicated variants were grown to logarithmic phase, and dropped on SG-Leu plates in a 1:5 dilution series. Plates were incubated at 30°C and imaged after 5 days. (B) WT or *yme1*Δ cells encoding the indicated Tgl2 variants were grown to mid-logarithmic phase and whole cell (W), cytosolic (C), or mitochondrial (M) fractions were isolated by mechanical lysis. The samples were analysed by SDS-PAGE and immunodecoration with antibodies against HA and Tom20 as a loading control. (C) Mitochondria isolated from either WT or *yme1*Δ cells expressing the indicated Tgl2 variants were solubilized with digitonin and analysed by BN-PAGE (4-14%) and immunodecoration with antibodies against HA or Tom40, as a loading control. (D) Aliquots (5%) of the solubilized mitochondria described in (C) were taken after the clarifying spin and analysed by SDS-PAGE and immunodecoration against HA and Tom20, as a loading control.

Next, we wanted to analyse if non-functional Tgl2 variants form higher molecular weight complexes, and if these complexes behave similar to the native one. We isolated mitochondria from *tgl2*Δ and *yme1*Δ cells expressing the variants of interest, solubilized them with digitonin, and analysed them by native PAGE and immunodecoration (Fig. 4C). We could detect the complex for HA-Tgl2 in mitochondria isolated from either both *tgl2*Δ and *yme1*Δ cells. In contrast, a complex containing Tgl2-HA could be detected only in mitochondria lacking Yme1 (Fig. 4C). Of note, HA-Tgl2_cys_ is unable to form a complex even in the absence of Yme1 indicating that the complex is dependent on the cysteine residues. As a control, the levels of the TOM complex, detected with an antibody against Tom40, are not altered (Fig. 4C). To confirm that the cells expressed the different proteins, a fraction the solubilized mitochondria was also loaded on SDS-PAGE and analysed using the indicated antibodies (Fig. 4D).

### Analysis of the contribution of single cysteine residues of Tgl2

The closest structural homologue of Tgl2 is a bacterial lipase – LipA, in the periplasmic space of *P. aeruginosa*. LipA has an intramolecular disulfide bond (between C183 and C235) which is important for the stability of the protein (28, 29). Disruption of this disulfide bridge results in loss of lipolytic function as well as rapid degradation of the protein (28). Since the Tgl2_cys_ variant is highly unstable (Fig. 4B), we wondered if Tgl2 has any specific intramolecular disulfide bonds that are essential for its stability. Using AlphaFold for structural prediction of Tgl2, we found cysteine residues (C31 and C150) that are potentially involved in intramolecular disulfide bridges (Fig. 5 and B, 30, 31). Additionally, we used bioinformatics tools (32–34) to predict the oxidation states of all the cysteine residues and to predict disulfide bridges between them (Fig. 5C). Based on these tools, we selected three cysteine residues – C31, C232, and C256, and replaced them by site-directed mutagenesis with alanine residues, creating HA-Tgl2_C31A_, HA-Tgl2_C232A_, and HA-Tgl2_C256A_, respectively. Next, we transformed them into *tgl2*Δ, *mcp2Δ/tgl2*Δ, and *yme1*Δ strains. Surprisingly, we observed that all three mutants could rescue the growth defect of *mcp2Δ/tgl2*Δ (Fig. 5D). Thus, it seems that despite their predicted participation in disulfide bridges, these cysteine residues are not essential for the functionality of the protein.

**Fig. 5.**
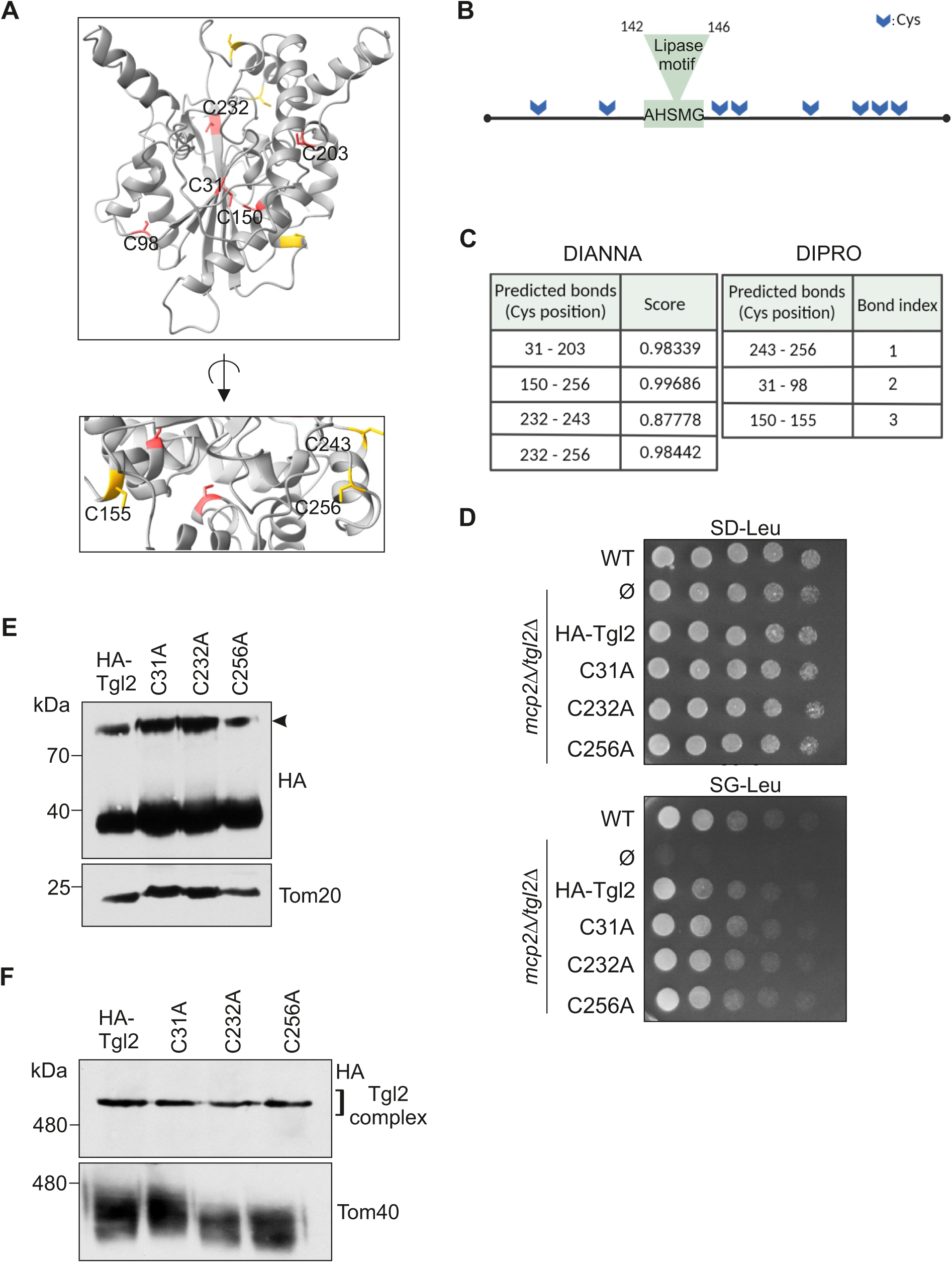
Analysis of the contribution of single cysteine residues of Tgl2. (A) Tgl2 has 8 cysteine residues that can potentially form intra or intermolecular disulfide bridges. Predicted structure of Tgl2 with the cysteine residues highlighted in red (functional group buried) or yellow (functional group exposed). (B) Schematic representation of the cysteine residues of Tgl2 marked with blue arrows and the canonical lipase motif marked in green (Created with Biorender). (C) Prediction of putative disulfide bonds within Tgl2 using using DiANNA and DIPro. The cysteine pairs are in decreasing order of probability of disulfide bond formation. (D) WT or *mcp2*Δ*/tgl2*Δ cells were transformed with an empty vector (Ø) or a plasmid encoding the indicated variants were grown to logarithmic phase, and dropped on SD-Leu or SG-Leu plates in a 1:5 dilution series. Plates were incubated at 30°C and imaged after several days. (E) *tgl2*Δ cells expressing the indicated variants of Tgl2 were grown to mid-logarithmic phase and the crude mitochondrial fractions were isolated. The samples were analysed by SDS-PAGE and immunodecoration with the indicated antibodies. (F) Mitochondria from *tgl2*Δ cells expressing the indicated protein variants were solubilized with digitonin and analysed by BN-PAGE (4-14%) and immunodecoration with antibodies against HA or Tom40.

To study the steady-state levels of the three mutants, we isolated crude mitochondrial fractions from *tgl2*Δ and *yme1*Δ yeast and analysed them by Western blotting. We observed that the expression levels and the stability of the cysteine variants were comparable to that of the native protein. Moreover, none of these point mutations had an impact on the homodimerization although C256 and C232 (partially) are exposed on the surface of the protein and could be involved in intermolecular disulfide bridges. (Fig. 5E). Since the Tgl2 complex is cysteine dependent (Fig. 4C), we wanted to analyse if residues C31, C232, and C256 are essential for its formation and/or stability. With that aim, we analysed isolated mitochondria harbouring the cysteine variants of Tgl2 by BN-PAGE (Fig. 5F). We could not observe any difference in the complex formation or stability between the native protein and the cysteine mutants. Thus, we conclude that these single point mutations by themselves are not sufficient to destabilize either the homodimer or oligomer formed by Tgl2 and therefore have no obvious impact on the functionality of the protein.

### The predicted catalytic triad of Tgl2 is required for its functionality

Tgl2 was first categorized as a lipase due to its canonical lipase motif - (G/A)XSXG, of which the serine is the catalytically active residue (Fig. 6A, adapted from (16)). In *in vitro* lipase assays using affinity tag-enriched protein, Ham et al reported that mutating the Serine 144 to Alanine (HA-Tgl2_S144A_) leads to loss of lipolytic activity of Tgl2 (14). Moreover, prokaryotic and eukaryotic lipases typically contain a catalytic triad consisting of Ser-Asp-His, wherein the serine is often part of the conserved (G/A)XSXG motif. This triad is also analogous to the catalytic centre of serine proteases as well as acyltransferases such as Tgl3 and Lro1 (35–37 and Fig. S3). We used structural data on LipA as well as other bacterial lipases that share homology with Tgl2, to gain more insight into the catalytic triad of Tgl2 (38). *In silico* analysis of the predicted structure of Tgl2 suggest a catalytic triad comprising of Ser144, Asp259, and His281 (Fig. 6B) (30, 31, 38).

**Fig. 6.**
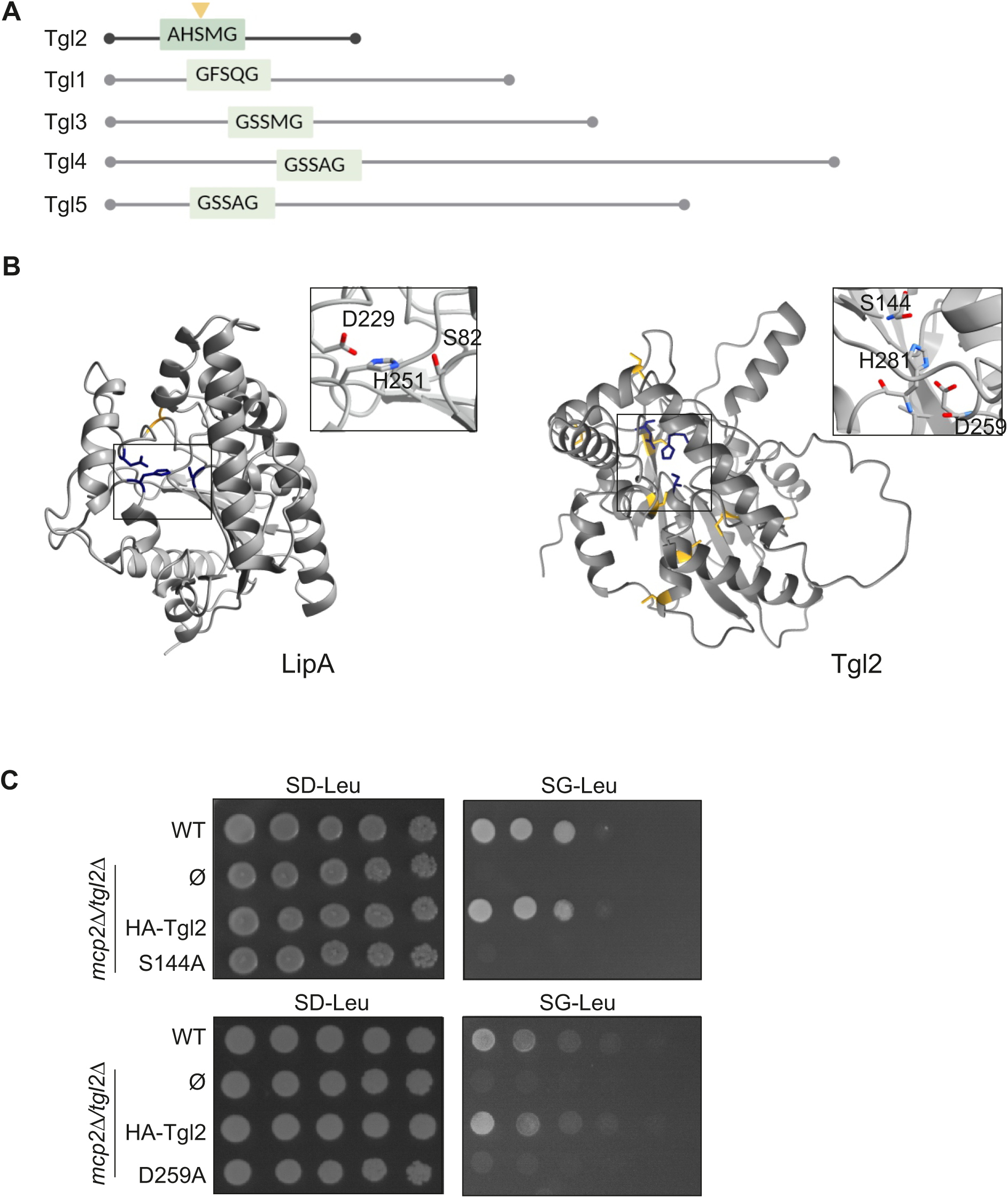
The predicted catalytic triad of Tgl2 is required for its functionality. (A) Tgl2 belongs to the Tgl family of yeast lipases and has a characteristic lipase motif (green) with a catalytically active serine (yellow arrow). Adapted from (16). (B) The predicted catalytic triad of Tgl2 (Ser-Asp-His). The crystal structure of LipA (left) with the intramolecular disulfide bond highlighted in yellow and the catalytic triad consisting of S82, D229, and H251 highlighted in blue ((29, 30) was employed to predict the structure of Tgl2 (right) with its cysteine residues in yellow and the predicted catalytic triad (S144-D259-H281) in blue. (C) WT or *mcp2*Δ*/tgl2*Δ cells transformed with the indicated variants were grown to logarithmic phase, and dropped on either SD-Leu or SG-Leu plates in a 1:5 dilution series. Plates were incubated at 30°C and imaged after several days.

We performed site-directed mutagenesis to mutate the catalytically active S144 (14) or D259 to alanine. Next, we tested the ability of the resulting HA-Tgl2_S144A_ and HA-Tgl2_D259A_ constructs to complement the growth phenotype exhibited by *mcp2Δ/tgl2*Δ cells. As expected, we could observe comparable growth behaviours of all the tested strains on fermentable carbon sources. In agreement with the previous reports, HA-Tgl2_S144A_ was unable to complement the growth phenotype exhibited by *mcp2Δ/tgl2*Δ on glycerol-containing medium (Fig. 6C). In addition, HA-Tgl2_D259A_ was also unable to rescue the growth phenotype under these conditions. These findings indicate that the residues S144 and D259 are potentially involved in the catalytic triad of the protein function.

### Disruption of the catalytic triad leads to protein instability

HA-Tgl2_S144A_ is a non-functional mutant which can be overexpressed to the same levels as the native protein and could also form the dimeric form of Tgl2 (Fig. 7A). To study the impact of this mutation on the quaternary structure of Tgl2, we solubilized mitochondria isolated from *tgl2*Δ and *yme1*Δ cell harbouring HA-Tgl2 or HA-Tgl2_S144A_ and subjected them to BN-PAGE and immunodecoration with antibodies against the HA-tag and Tom40.

**Fig. 7.**
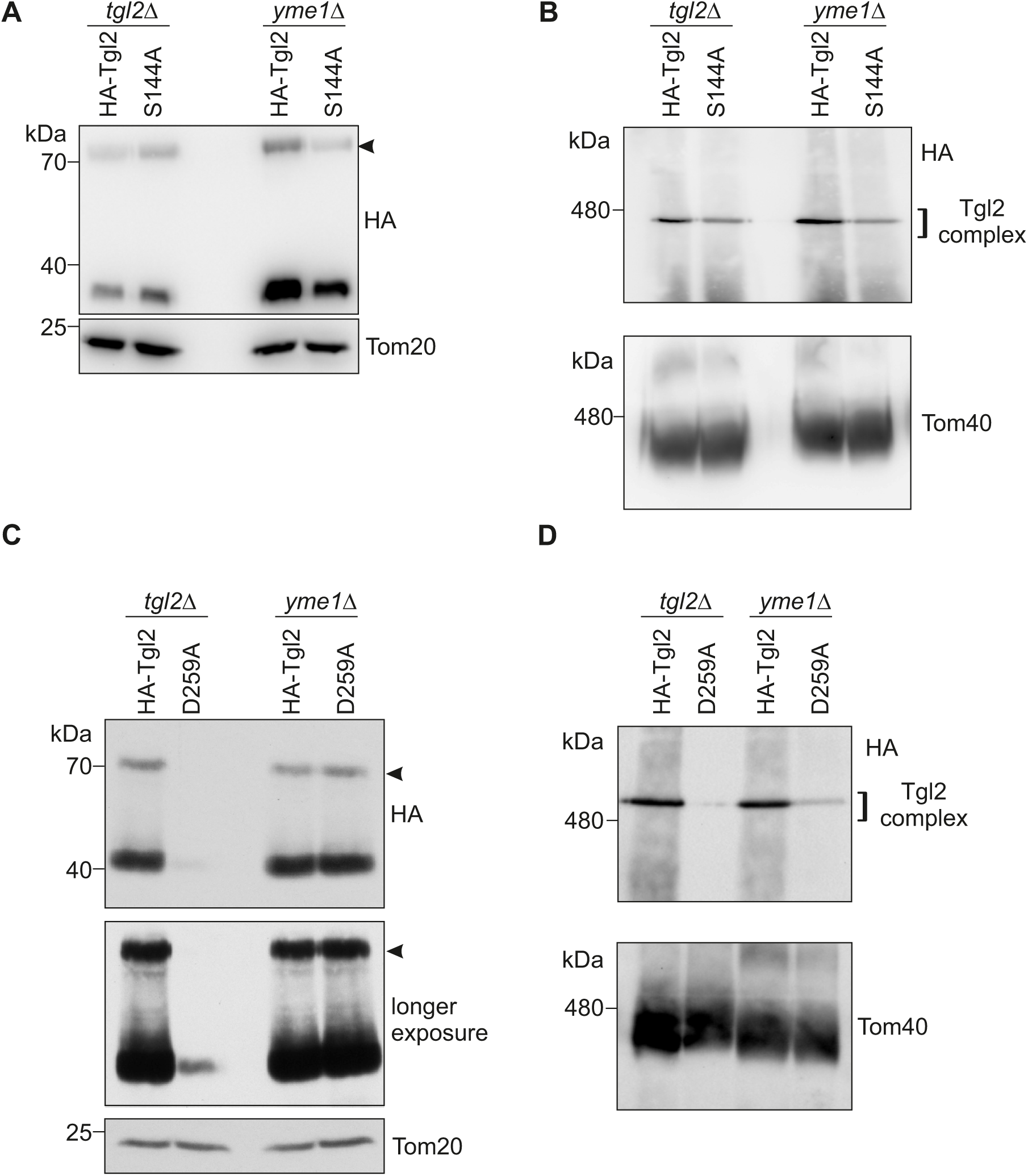
Disruption of the catalytic triad leads to protein instability. (A) and (C) WT and *yme1*Δ cells expressing the indicated Tgl2 variants were grown to mid-logarithmic phase and crude mitochondrial fractions were isolated by mechanical lysis. The samples were analysed by SDS-PAGE and immunodecoration with antibodies against HA and Tom20, as a loading control. (B) and (D) Mitochondria isolated from WT and *yme1*Δ cells expressing the indicated Tgl2 variants were solubilized with digitonin and analysed by BN-PAGE (4-14%) and immunodecoration with antibodies against HA and Tom40.

Figure 7B shows that the complex could be detected comparably for both variants. We conclude that while the Ser144 is vital for the catalytic activity of the protein, it has no impact on the stability or structural integrity of the monomeric, dimeric, or oligomeric form of the protein.

To analyse if the catalytically active aspartate is crucial for protein stability, we analysed its steady-state levels in *tgl2*Δ and *yme1*Δ yeast cells. We found that in addition to losing its function, the aspartate mutant is expressed notably less than the native protein in *tgl2*Δ cells yet to the same extent in *yme1*Δ (Fig. 7C). This expression behaviour was similar for the protein complex, as the higher molecular weight complex is detectable only in very little amounts for the mutated protein in *tgl2*Δ cells, but rather similar to the native complex in *yme1*Δ yeast (Fig. 7D). In contrast to the S144A variant, perturbing the catalytic triad in mutating the aspartate residue leads to instability in complex formation in WT-like conditions, suggesting that the aspartate residue is critical for the stability of Tgl2.

These observations are in congruence with previous studies where the catalytically active aspartate residue was shown to be crucial for protein stability in lipases from *G. candidum* as well as human pancreatic lipase and choline esterase. These previous studies have also reported that while mutating the serine residues in the catalytic triad leads to loss of function, it has no impact on protein stability (39).

### Genetic interactions of *MCP2*, *TGL2,* and *YME1*

In this study, we assess several different mutants of Tgl2 to gain a better understanding of the protein’s structure-function relationship. We use complementation of *mcp2/Δtgl2*Δ under non-fermentable conditions as a tool to analyse functionality. Of the mutants created in this study, the non-functional variants (HA-Tgl2_Cys_, Tgl2-HA, HA-Tgl2_S144A,_ and HA-Tgl2_D259A_) either could not be detected or were detected in very low amounts in *mcp2Δ/tgl2*Δ (Fig. S2). Naturally, this raises the question – are these variants indeed non-functional due to loss of enzymatic capacity or are they unable to complement the growth phenotype simply due to their low expression levels.

Since we could show that Yme1 is responsible for the turnover of Tgl2 (Fig. 4B), we attempted to create the triple mutant strain - *mcp2Δ/tgl2Δ/yme1*Δ to ensure that the mutated Tgl2 variants are not degraded and the growth defect caused by the absence of Mcp2 and Tgl2, can be analysed in the absence of Yme1. Interestingly, the deletion of all three genes always yielded non-viable spores. Figure 8A shows an excerpt from n>100 tetrads dissected.

**Fig. 8.**
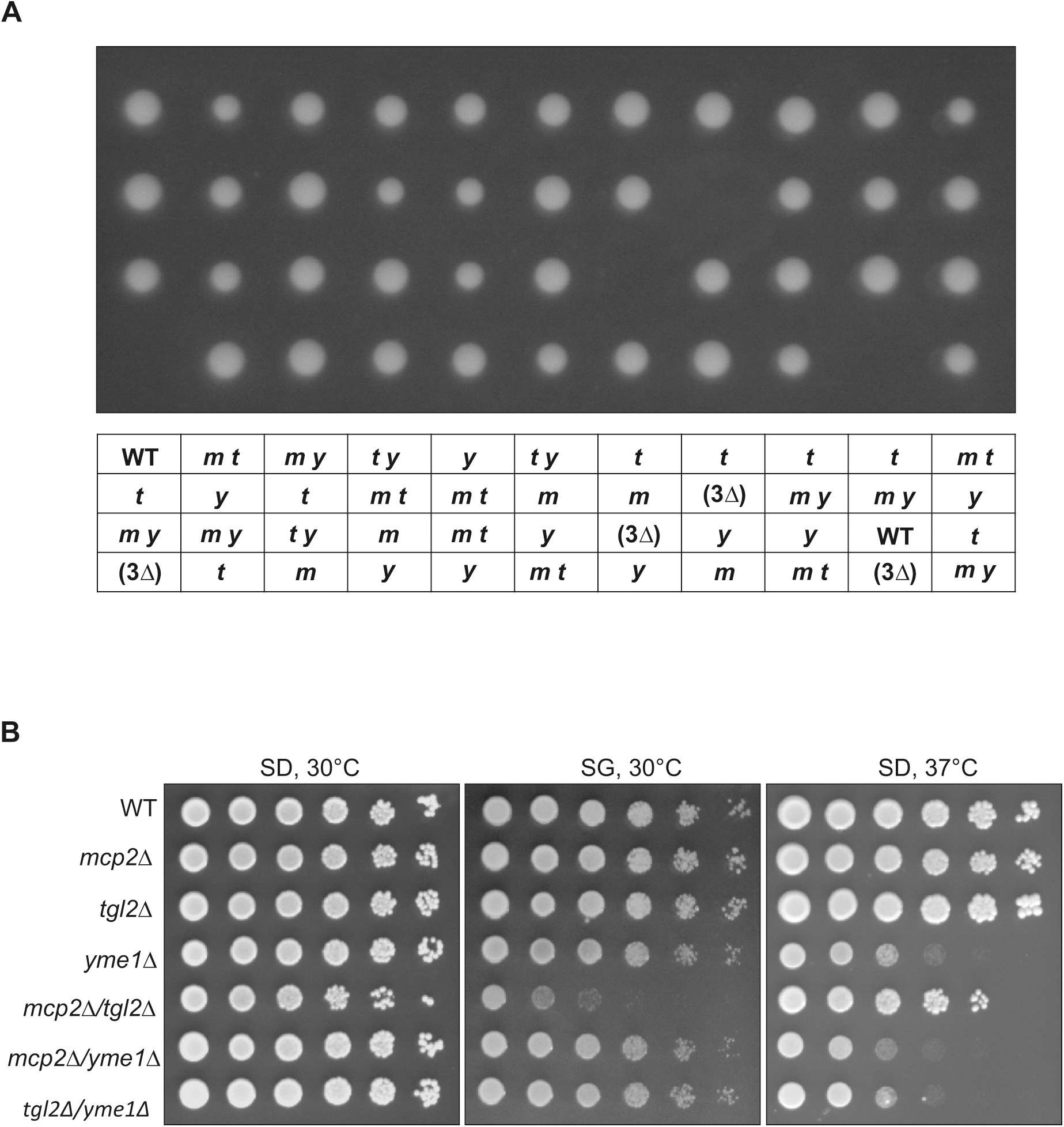
Genetic interactions of *MCP2*, *TGL2,* and *YME1*. (A) Excerpt of 11 out of more than 100 dissected tetrads of the heterozygous triple deletion mutant (*mcp2Δ/MCP2 - tgl2Δ/TGL2 - yme1Δ/YME1*). The table below the haploid clones shows the genotype analysis, *m*: *mcp2Δ*, *t*: *tgl2Δ*, *y: yme1Δ,* WT: wild type, (3*Δ*): triple deletion – not viable. (B) Growth analysis of single and double deletion strains of the three genes. Growth at either 30°C or 37°C was analysed by drop dilution assay of 1:5 serial dilutions on synthetic medium containing either glucose (SD) or glycerol (SG).

Of note, we observed that lethality of the triple deletion mutant does not stem from a simple combination of growth defects caused by the three individual genes. All three single deletion mutant cells as well as all combinations of double mutations show no growth defect on glucose containing synthetic medium at normal temperatures. Furthermore, the growth defect of *yme1*Δ cells at elevated temperatures is not worsened by loss of Mcp2 or Tgl2. Finally, on non-fermentable carbon sources where *mcp2*Δ/*tgl2*Δ cells show the pronounced growth defect, *yme1*Δ cells grow like wild type (Fig. 8B).

Although we so far cannot explain the cause of lethality of the combined deletion of *MCP2*, *TGL2* and *YME1,* we assume that all three proteins share a functional relationship. As expected, when we express functional plasmid borne Tgl2 in heterozygous diploid triple mutant cells prior to tetrad dissection we can retrieve triple deletion cells expressing plasmid borne Tgl2. On the other hand, empty plasmids or plasmids encoding the non-functional Tgl2 variant Ser144A failed to yield triple deletion cells expressing the mutant proteins (data not shown).

### Mitochondria lacking Tgl2 have elevated levels of neutral lipids

Previous lipidomics analysis from whole cell extracts, indicated elevated TAG and DAG levels in *mcp2Δ/tgl2*Δ yeast cells (15). Since Tgl2 and Mcp2 are mitochondrial proteins, we wondered if this perturbation is primarily caused by mitochondrial alternations. To that goal, we extracted neutral lipids from isolated mitochondria (WT, *mcp2*Δ, *tgl2*Δ, and *mcp2Δ/tgl2*Δ) and subjected them to thin-layer chromatography (TLC) analysis. We observed a slight increase in TAG levels in organelles from *mcp2Δ/tgl2*Δ (Fig. 9A). To further analyse the lipid composition of the isolated organelles, we trypsinized the mitochondria from WT, *mcp2*Δ, *tgl2*Δ, and *mcp2Δ/tgl2*Δ, to eliminate protein-mediated mitochondrial contact sites, and purified the organelles by a density gradient. The purity of the mitochondria was confirmed by SDS-PAGE and immunodecoration with antibodies against peroxisomal (Pex14), ER (Erv2), and mitochondrial (Tom40 and Hep1) marker proteins. Figure 9B confirms that the mitochondrial fractions are essentially free from ER and/or peroxisome contaminations. Lipidomics analyses of the purified mitochondria revealed elevated TAG and DAG levels in *mcp2Δ/tgl2*Δ (Fig. 9C). Collectively, these results suggest that the combined deletion of TGL2 and MCP2 results in an increase mitochondrial or mitochondria-associated TAGs.

**Fig. 9.**
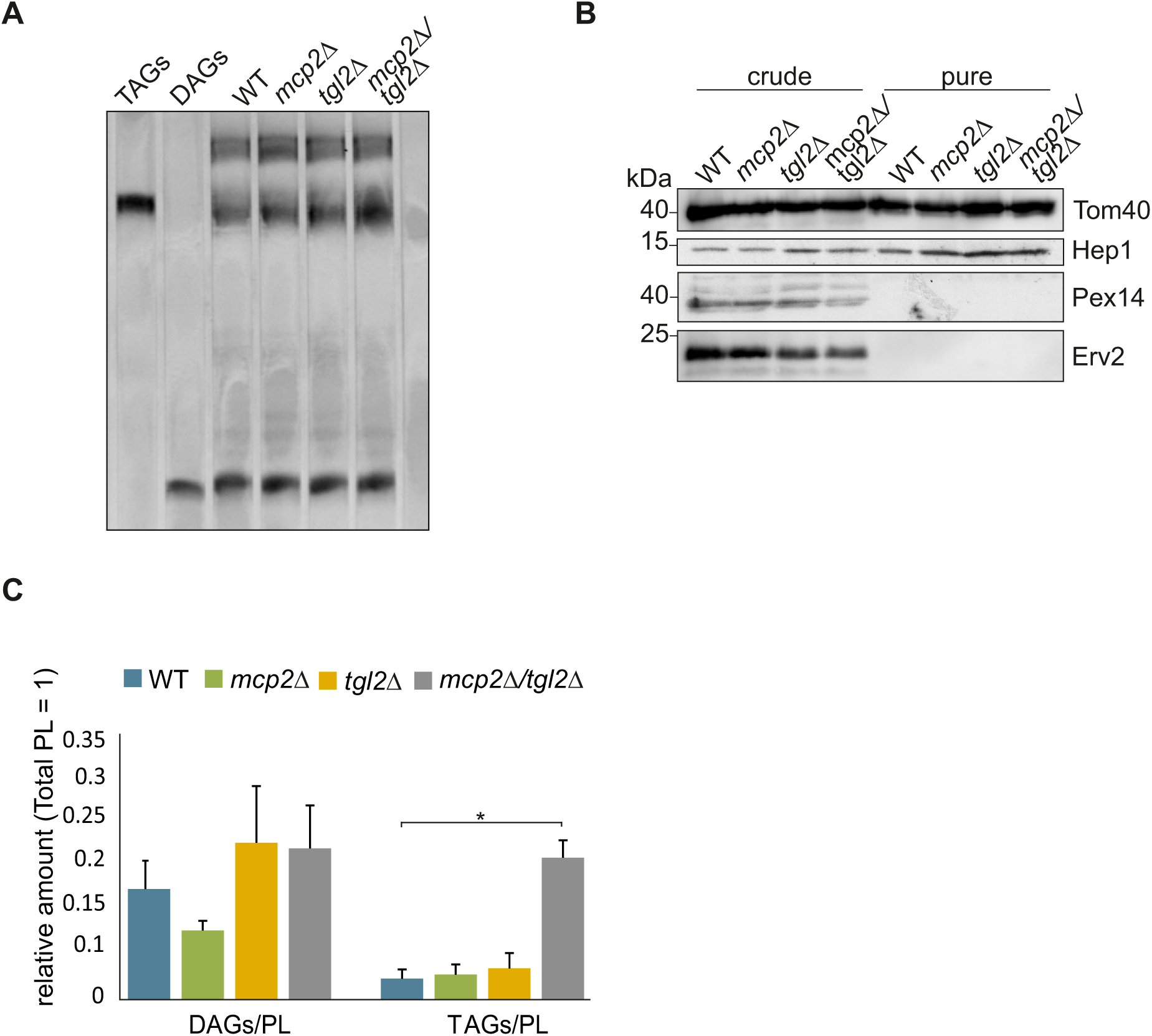
Mitochondrial fractions from *mcp2*Δ/*tgl2*Δ cells show increased TAG levels. (A) Neutral lipids were extracted from mitochondria of the indicated cells and analysed by thin-layer chromatography. The lipids were stained with a primuline solution and visualised under UV-light. The migration behaviour of standard neutral lipids is indicated on the left. (B) Mitochondria were isolated from the indicated strains, treated with trypsin and further purified by sucrose gradient. Crude and purified mitochondria (100 μg) were analysed by SDS-PAGE and immunodecoration. Tom40 and Hep1, mitochondrial markers, Pex14, peroxisomal marker, and Erv2, ER marker. (C) Pure mitochondria from (B) were analysed by mass spectrometric analysis. The amounts of TAGs and DAGs versus total PL is shown as a mean with SD bars (n = 3, * p < 0.05, GraphPad T-test calculator).

## Discussion

The importance of cellular lipid homeostasis is highlighted by its implication for human health (40, 41). Even in simpler model organisms like *S. cerevisiae*, deficiencies in lipid metabolism result in impaired cellular function and, in some cases, severe growth defects (40, 41). There have been significant advancements in the field of lipid metabolism in the last few years wherein the cross talk between mitochondria and other organelles has acquired a crucial role. Reports of protein tethers between the mitochondrial outer membrane (MOM) and membranes of proximal organelles laid the foundation of the concept of ‘contact sites’, which has become a topic of extensive research. The ERMES complex was the first of such mitochondrial tethers to be recognized, and has been now thoroughly elucidated (42–44). ERMES subunits have been shown to bind to and traffic phospholipids between the ER and mitochondria. The loss of these subunits results in severe growth phenotypes and anomalous mitochondrial morphology (45–48).

In a previous study, we characterized a high copy suppressor of ERMES mutants – Mcp2 (now Cqd2) which was more recently suggested to be involved in CoQ metabolism in yeast (19). To better characterize *MCP2/CQD2*, we performed a high throughput screen of *mcp2Δ* and identified *TGL2* as a hit, where the two genes displayed negative genetic interaction, more prominently on non-fermentable carbon sources (15). This growth defect is lost after prolonged growth on fermentable carbon sources – a phenomenon often observed for lipid metabolism defects and ERMES mutants (46, 49, 50). We could also show that both Mcp2 and Tgl2 are located with their functional domains in the IMS. Of the two, Tgl2 is a soluble protein and relies on the disulfide relay pathway for its import into the IMS (15).

In our current study, we further characterized Tgl2 to gain a better understanding of its molecular structure and function. We observed that Tgl2 can form homodimers under non-reducing conditions and suggest that the dimer is a product of intermolecular disulfide bridges between one of the eight cysteine residues of each monomer of the protein. We could also confirm an oligomeric form of the protein containing at least two copies of Tgl2. This complex is also thiol dependent and is susceptible to treatment with stronger detergents. The formation of a structural important disulfide bridge would agree with a more oxidising environment in the IMS, similar to the periplasm of bacteria. Moreover, Tgl2 is not conserved amongst higher eukaryotes and has its closest structural homologues in prokaryotes. These bacterial lipases e.g. LipA from *P. aeruginosa*, were shown to harbour intra- as well as intermolecular disulfide bridges. Furthermore, extracellular lipases of higher eukaryotes like human pancreatic lipase, and lipoprotein lipase also contain disulfide bonds which are often essential for protein function, allowing dimerization that facilitates substrate binding or protein stability (28, 51, 52). Since mutation of all cysteine residues of Tgl2 led to loss of function of the protein we started to mutate three single cysteine residues that according to structure prediction programs are likely to be involved in intra- or intermolecular cysteine bridges. However, replacing the cysteines 31, 232, or 256 to alanine had no influence on the quaternary structure or function of Tgl2. We cannot exclude the possibility that the function of the cysteine residues is somewhat redundant, and therefore a single replacement can be compensated by the other seven cysteine residues.

The non-functional variants of Tgl2 like Tgl2-HA and Tgl2_cys_, are nearly undetectable in WT or *tgl2Δ* cells despite their over-expression, indicating a strict quality control mechanism for the protein. A vast majority of the IMS proteins rely on Yme1, an i-AAA protease in the MIM, for their folding and/or degradation (26). Yme1 has an IMS exposed catalytic domain which is suggested to have chaperone-like activity (27). By comparing the steady state levels of native and non-functional variants of Tgl2, we could confirm that its quality control is maintained by Yme1. The undetectable or weakly detectable protein mutants could be observed upon deletion of *YME1*, in amounts similar to those of native Tgl2. We also found a synthetic lethal genetic interaction of *MCP2*, *TGL2*, and *YME1* which suggests a functional relationship beyond the quality control of Tgl2 by Yme1. Further studies are required to understand this observation.

Ham et al. previously characterized the lipase motif of Tgl2. By mutating the conserved serine residue (S144), they could show a loss of *in vitro* lipolytic activity (14). Our observations that despite native-like expression levels, Tgl2_S144A_ is unable to rescue the growth defect of *mcp2*Δ*/tgl2*Δ, agree with their report. We also characterized the predicted Ser-Asp-His catalytic triad of Tgl2 and showed that disrupting the catalytically active triad by replacing the aspartate by alanine leads to an unstable, poorly expressing variant of the protein (51). Studies pertaining to other lipases and acyltransferases show that while a conservative mutation (aspartate to glutamate) results in only partial loss of function, mutating the aspartate to alanine or valine leads to loss of enzyme activity and stability. Since the carboxylic group is essential for stabilizing interactions with other residues in its vicinity, its loss can lead to protein misfolding and/or instability (37, 39)

The catalytic triad is a feature of several enzymes including proteases, acyltransferases, and lipases (Fig. S3 and (51)). Some members of the Tgl family exhibit lipase as well as acyltransferase activity, therefore Tgl2 could also be involved in transferring fatty acids between substrates rather than hydrolysing TAGs like classical lipases (16, 53).

The lipolytic activity of Tgl2 was shown through *in vitro* lipase assays wherein Tgl2 enriched lysate was used to hydrolyse different TAGs – of which tributyrin was the optimal substrate (14). Of note, tributyrin is not a typical physiological substrate in mitochondria. We performed lipidomics analysis of purified mitochondrial fractions and could detect elevated levels of neutral lipids in fractions isolated from *mcp2*Δ/*tgl2*Δ yeast cells. This observation aligns with our previous lipidomics analysis from whole cell lysate fractions of the same cells (15). Notably, the neutral lipid content measured in the lipidomics analyses reflected primarily long chain fatty acids, which are the most abundant forms of TAGs and DAGs in yeast cells. In conclusion, our experiments could show that the combined absence of Tgl2 and Mcp2 results in increased neutral lipid amounts in yeast cells which is mainly caused by an increase in TAGs in mitochondrial fractions. Further experiments are required to clarify whether TAGs indeed occur in between the leaflets of either the MIM or the MOM. We cannot exclude that stronger interaction between LDs and mitochondria, in the absence of Mcp2 and Tgl2, leads to an increased LD co-sedimentation during isolation of mitochondria. The physiological function of Tgl2 in the IMS is currently unclear and further studies are required to confirm whether it behaves as a classical TAG lipase or it evolved to perform a different role. To achieve a comprehensive understanding of Tgl2’s function, we tried to identify its physical partners through co-immunoprecipitation experiments followed by mass spectrometry experiments. Despite using different affinity tags and experimental conditions, we were unable to find any proteins of interest in these assays. In a recent mitochondrial complexome study, Tgl2 was shown to have a similar migration behaviour as Ups1 and Ups3, lipid transfer proteins in the IMS (25). To investigate potential Tgl2-Ups1 interactions, we performed BN-PAGE with *ups1*Δ yeast cells expressing HA-Tgl2 but the complex remains unchanged in size and stability (Fig. S1). We applied the same approach to *ups2*Δ and *mdm35*Δ, proteins involved in the lipid transfer complex of the IMS (22, 23), but the complex was still unchanged (Fig. S1).

The closest structural homologue of Tgl2 is a bacterial TAG lipase – LipA, which is located in the periplasm. Given the bacterial ancestry of mitochondria and the similarity of Tgl2 with mostly bacterial lipases, one could speculate that during the course of evolution, the protein was maintained in simpler eukaryotes like yeast but was lost in higher organisms. Taken together, our study sheds new light on the structure-function relationship of the mitochondrial protein Tgl2.

### Experimental procedures

#### Yeast strains and growth conditions

Yeast strains were grown in standard rich medium (YP) or synthetic medium (S) with either glucose, galactose, or glycerol as the carbon source.

Transformation of yeast cells was done using the lithium acetate method. For strains transformed with plasmids, synthetic media with appropriate selection marker(s) were used.

For drop dilution assays, yeast cells were cultured to an OD_600_ of 1.0 and serially diluted in five-fold increments. An aliquot of the serially diluted cultures (5 µL) was spotted onto the corresponding solid medium, and the cells were incubated at 30 or 37°C.

All deletion strains were made using tetrad dissection and confirmed by PCR using primers specific for the genes of interest.

All *S. cerevisiae* strains used in this study are listed in Table 2.

#### Recombinant DNA techniques

The previously constructed plasmid pGEM-Tgl2 (15) was used as a template for inserting a FLAG tag at the N-terminus of the ORF of Tgl2. The construct pGEM-HATgl2 was created using primers from (15) and this construct served as a template for site-directed mutagenesis reactions. Primer pairs containing the mutation of interest were used to generate a mutated sequence of Tgl2. The PCR products were digested using DpnI and transformed into *E. coli* cells for further screening. After confirmation with sequencing, the mutated constructs were cloned into a yeast expression plasmid.

All primers and plasmids used in this study are listed in Table 1 and 3, respectively.

**Table 1:**
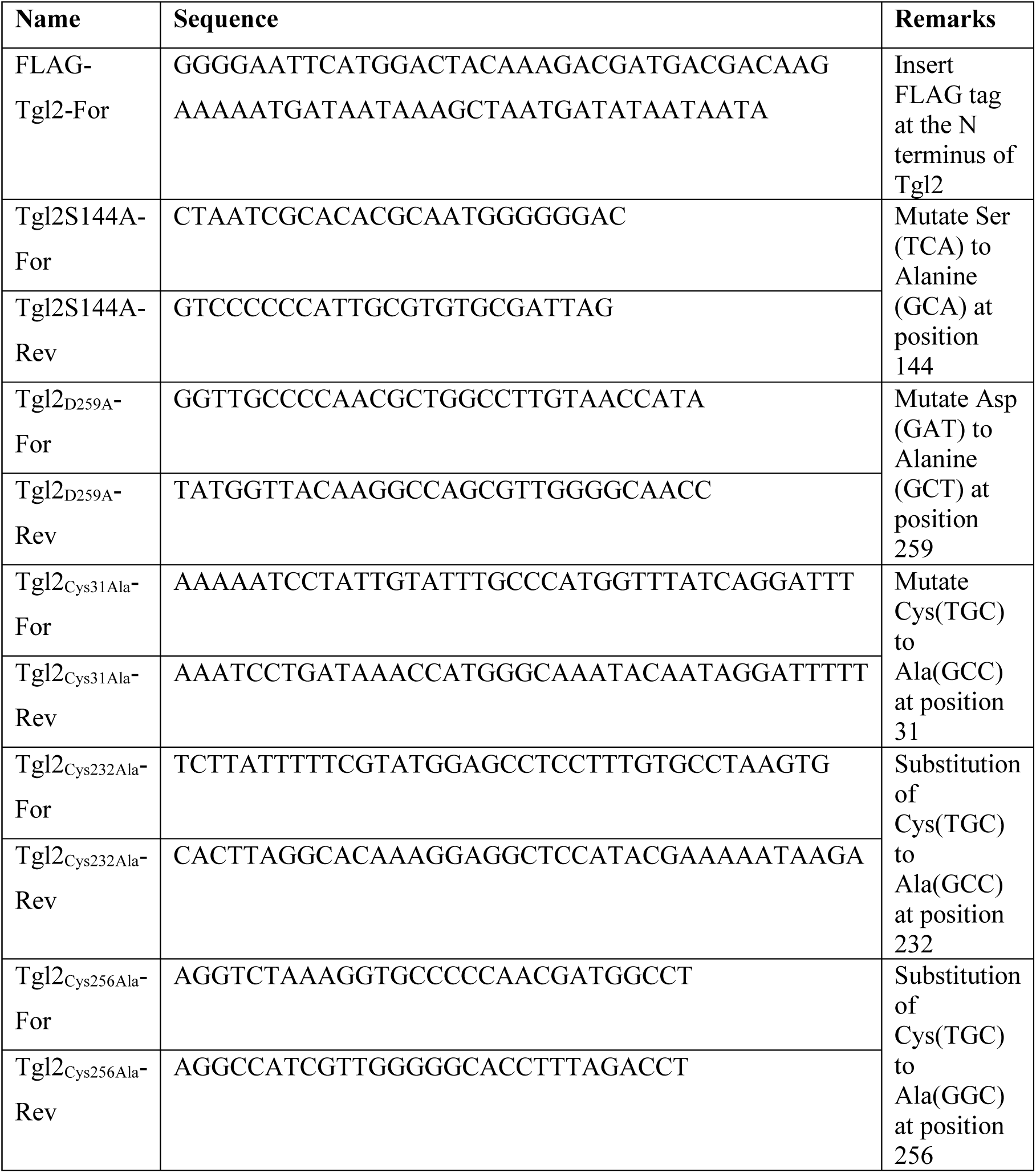
Primers used in this study.

**Table 2:**
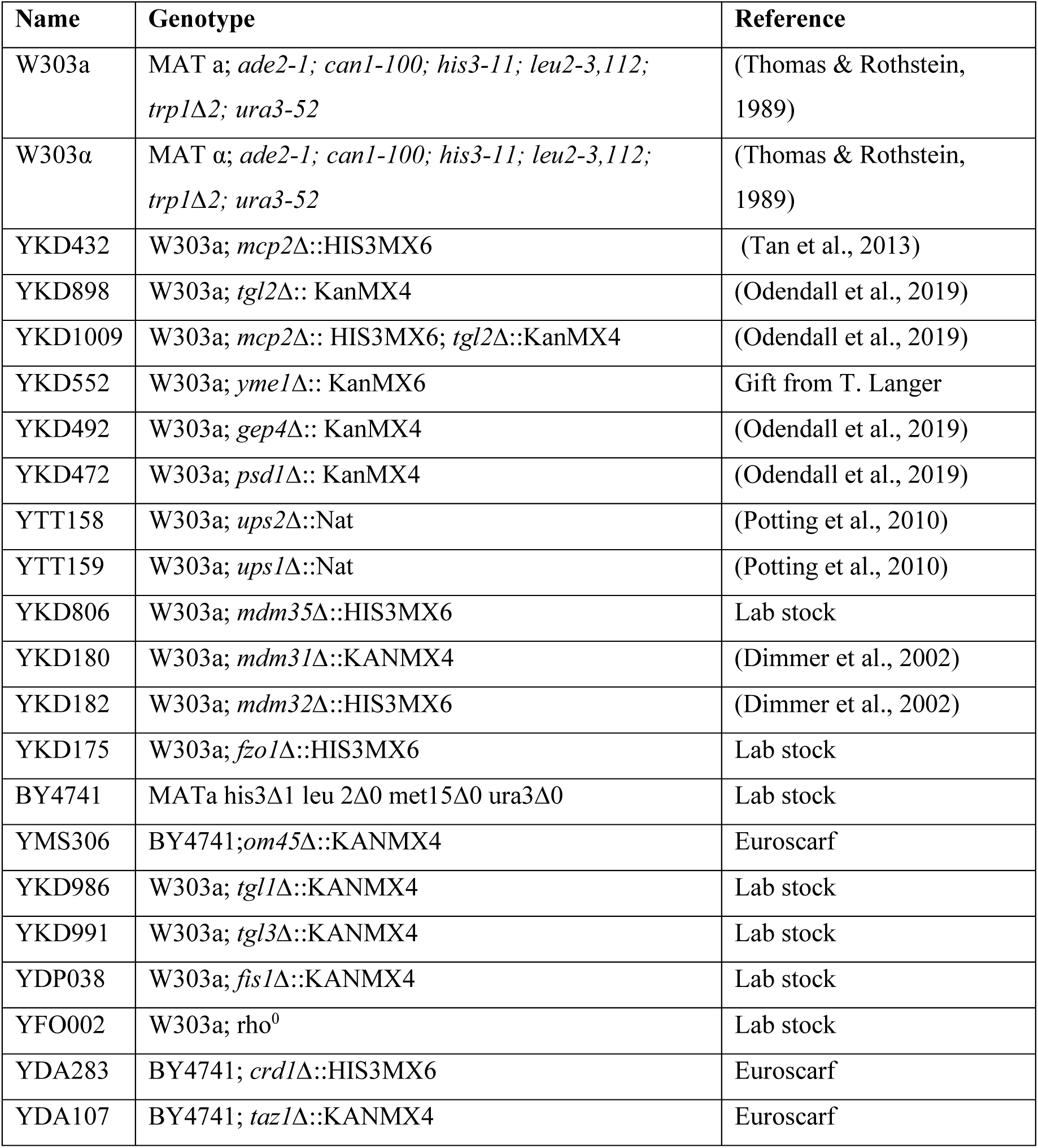
*S. cerevisiae* strains used in this study.

**Table 3:** List of plasmids used in this study.

| Name | Promoter | Markers | Coding Sequence |
| --- | --- | --- | --- |
| pYX142-HA-Tgl2 | TPI | LEU2,<br>Amp<sup>R</sup> | Tgl2 with N-terminal HA-Tag |
| pYX142-Tgl2-HA | TPI | LEU2,<br>Amp<sup>R</sup> | Tgl2 with C-terminal HA-Tag |
| pYX142-HA-Tgl2<sub>Cys</sub> | TPI | LEU2,<br>Amp<sup>R</sup> | Tgl2 with N-terminal HA-Tag, all cysteines mutated to serine |
| pGEM4-HA-Tgl2 | T7 | Amp<sup>R</sup> | Tgl2 with N-terminal HA-Tag |
| pYX122-FLAG-Tgl2 | TPI | HIS3,<br>Amp<sup>R</sup> | Tgl2 with N-terminal FLAG-Tag |
| pYX142-HA-Tgl2<sub>S144A</sub> | TPI | LEU2,<br>Amp<sup>R</sup> | Tgl2 with N-terminal HA-Tag, S144A mutation |
| pYX142-HA-Tgl2<sub>C31A</sub> | TPI | LEU2,<br>Amp<sup>R</sup> | Tgl2 with N-terminal HA-Tag, C31A mutation |
| pYX142-HA-Tgl2<sub>C232A</sub> | TPI | LEU2,<br>Amp<sup>R</sup> | Tgl2 with N-terminal HA-Tag, C232A mutation |
| pYX142-HA-Tgl2<sub>C256A</sub> | TPI | LEU2,<br>Amp<sup>R</sup> | Tgl2 with N-terminal HA-Tag, C256A mutation |
| pYX142-HA-Tgl2<sub>D259A</sub> | TPI | LEU2,<br>Amp<sup>R</sup> | Tgl2 with N-terminal HA-Tag, D259A mutation |

#### Isolation of mitochondria

Mitochondria were isolated from yeast cells using a previously published protocol based on differential centrifugation (54). Yeast cultures (2-5 L) were grown at 30°C and harvested at an OD_600_ of 0.8-1.6 by centrifugation (3000 xg, 5 min, RT). The cell pellets were washed, weighed and resuspended in resuspension buffer (100 mM Tris and 10 mM DTT). The cell suspension was incubated at 30°C for 10 min after which the cells were harvested as above. The cell pellets were washed with a spheroplasting buffer (1.2 M Sorbitol and 20 mM Sodium Phosphate, pH 7.2). Next, the cell pellets were resuspended in zymolyase-containing spheroplasting buffer and incubated at 30°C for at least 1 h with constant shaking. The spheroplasts were harvested (1100 xg, 5 min, 2°C), resuspended in Homogenization buffer (0.6 M Sorbitol, 1 mM EDTA, 1 mM PMSF, 0.2% (w/v) fatty-acid free BSA, and 10 mM Tris pH 7.4), and dounced 10-12 times. The lysed spheroplasts were first subjected to a clarifying step (2000 xg, 5 min, 2°C) and then to a high-speed centrifugation step (17,500 xg, 15 min, 2°C). The pellets were washed with the isotonic SEM buffer (250 mM sucrose, 1 mM EDTA, and 10 mM MOPS/KOH pH 7.2) containing 2 mM PMSF and centrifuged as in the previous step. The pellet obtained after the high-speed spin is the mitochondrial fraction. The pellet was resuspended in SEM buffer, snap-frozen in liquid N_2_, and stored at −80°C until further use.

To obtain pure mitochondrial fractions, the mitochondrial pellet obtained after differential centrifugation was subjected to trypsinization (5 µg of trypsin per mg of mitochondrial protein) on ice for 20 min. The reaction was stopped by adding Soybean Trypsin Inhibitor (STI) (10 µg STI per µg of Trypsin) and incubated on ice for 30 min. The trypsinized mitochondria were mixed with 2-3 mL of SEM buffer containing 1 mM PMSF and loaded on a sucrose step gradient (20, 30, 40, 50, and 60% (w/v) sucrose in 10 mM MOPS/KOH pH 7.4, 100 mM KCl, 1 mM EDTA, 1 mM PMSF). The gradients were centrifuged (210,000 xg, 16 h at 4°C) using a swing-out rotor (SW40Ti). The fractions between the 40%-60% phase - the mitochondrial fractions, were collected and diluted with 35 mL SEM buffer containing 2 mM PMSF. A pure mitochondrial pellet was harvested by centrifuging the suspension (17500 xg, 15 min at 4°C). The pellet was resuspended in SEM buffer, snap-frozen, and stored at −80°C until further use.

#### Alkaline Extraction

Isolated mitochondria (50 µg) were resuspended in 50 µL of 20 mM MOPS/KPH pH 7.5. Next, 50 µL of 0.1 M Sodium carbonate solution of varying pH (11 or 11.5) was added to the samples, which were then incubated on ice for 30 min. The samples were centrifuged (94000 xg, 30 min, 4°C), the pellet was resuspended in sample buffer. The supernatant was subjected to trichloroacetic acid (TCA) precipitation and the resulting pellet was also resuspended in sample buffer. The samples were boiled at 95°C for 10 min and analysed by SDS-PAGE and immunodecoration.

All antibodies used in this study are listed in Table 4.

**Table 4:** List of antibodies used in this study.

| Antibody | Dilution | Source |
| --- | --- | --- |
| Polyclonal rat anti-HA | 1:1000 | Roche |
| Polyclonal rabbit anti-Erv2 | 1:1000 | Lab of Roland Lill |
| Polyclonal rabbit anti-Hep1 | 1:2000 | Lab stocks |
| Polyclonal rabbit anti-Pex14 | 1:10000 | Lab stocks |
| Polyclonal rabbit anti-Bmh1 | 1:10,000 | Lab stocks |
| Polyclonal rabbit anti-Tom20 | 1:4000 | Lab stocks |
| Polyclonal rabbit anti-Tom40 | 1:4000 | Lab stocks |
| Polyclonal rabbit anti-Tom70 | 1:2000 | Lab stocks |
| Polyclonal rabbit anti-FLAG | 1:2000 | ThermoFischer |
| Horseradish peroxidase coupled goat anti-rabbit | 1:10000 | 1721019 (Bio-rad) |
| Horseradish peroxidase coupled goat anti-rat | 1:5000 | ab6845 (Abcam) |

#### Isolation of mitochondria by mechanical rupture

Yeast cultures (100-200mL) were grown at 30°C and harvested at an OD_600_ of 0.8-1.6 by centrifugation (3000 xg, 5 min, RT). The pellet was resuspended in SEM buffer and the cells were disrupted by repeated cycles of vortexing with glass beads at 4°C. The lysed cells were centrifuged (2000 xg, 3 min, 4°C) to remove unbroken cells and the crude mitochondria were pelleted from the supernatant by centrifugation (14000 xg, 12 min 4°C).

#### Co-Immunoprecipitation

Mitochondria (1 mg) were re-isolated and solubilized in 500 µL of Tris-buffered saline (TBS) supplemented with protease inhibitor cocktail and 1% (w/v) digitonin. The mitochondria were incubated on an orbital shaker for 30 min at 4°C and the solubilized material was centrifuged (18000 xg, 30 min, 4°C). Magnetic Anti-FLAG beads were incubated with the supernatant on an orbital shaker for 1 h at 4°C. The beads were washed three times with 500 µL of TBS supplemented with protease inhibitor cocktail, and the bound proteins were eluted using a three-fold excess of FLAG peptide (150 ng/µL in TBS).

#### BN-PAGE

Mitochondria (50 µg) were solubilized in 50 µL of SEM buffer supplemented with either Triton X-100 (0.5% (v/v)) or digitonin (1% (w/v)). The solution was incubated for 30 min at 4°C on an orbital shaker and the solubilized fraction was subjected to a clarifying spin (20000 xg, 15 min, 4°C). The supernatant was mixed with the loading dye (5% (w/v) Coomassie blue G, 500 mM 6-amino-N-caproic acid, 100 mM Bis-Tris, pH 7.0) and analysed on a 4-14% acrylamide blue native gel. The gels were blotted onto a PVDF membrane and further analysed by immunodecoration.

#### Antibody shift assay

Mitochondria (50 µg) were resuspended in 50 µL of SEM buffer supplemented with 1% digitonin and incubated on an orbital shaker for 30 min at 4°C. The solubilized mitochondria were subjected to a clarifying spin (20000 xg, 30 min, 4°C) and the supernatant was incubated with 1 µL of SEM buffer or 0.5 µL or 1 µL of anti-HA antibody. The samples were incubated on ice for 30 min and then centrifuged briefly (13000 xg, 10 min, 4°C). The supernatant was mixed with the loading dye as above and analysed by BN-PAGE and immunodecoration.

#### Extraction and analysis of mitochondrial neutral lipids

Lipid extraction of isolated mitochondria was done according to a published protocol (55).

Briefly, 500 µg of mitochondria were resuspended in a mixture of chloroform:methanol 1:2 (v/v), vortexed, and incubated on ice for 30 min. Next, 1.25 volumes of chloroform followed by 1.25 volumes of water were added - the samples were vortexed vigorously after each addition. The organic and inorganic phases were separated by centrifuging the samples (1000 xg, 10 min, RT). The lower, organic phase was collected with a Pasteur pipette and the samples were evaporated by SpeedVac. The lipids were resuspended in 30 µL of chloroform:methanol 1:1 (v/v) and immediately used for TLC.

Neutral lipids were separated by TLC using glass plates of pre-coated 0.25 mm silica gel with fluorescent indicator UV_254_. The silica plates were prepared by washing them with a mixture of chloroform: methanol 1:1 (v/v) and then allowing them to dry for 15-20 min. The plates were then sprayed with a solution of 2.3% boric acid (w/v) in ethanol and dried in an oven for 15 min at 100°C.

The lipids were spotted gradually (3-5 µl at a time) until the entire volume was spotted onto the TLC plate. The spots were air dried and then separated by a solvent mixture of chloroform/ethanol/water/trimethylamine (30:35:7:35, v/v/v/v). After the samples have migrated through the whole distance completely, the plate was air dried and the run was repeated to ensure successful separation. The plate was allowed to dry completely and then sprayed with a 0.05% (w/v) primuline solution in 80% (v/v) acetone. The plate was visualised under a UV-lamp.

#### Sucrose Gradient

A sucrose gradient (0.9/1/1.1/1.2/1.3 M) to separate MIM and MOM vesicles was performed according to a previously published protocol (18). Briefly, Mitochondria (3 mg) were reisolated and resuspended in 10 mL of swelling buffer (20 mM HEPES/KOH pH 7.4, 2 mM EDTA, 2 mM PMSF) and incubated for 1 h at 4°C on an orbital shaker. The sucrose concentration was adjusted to 0.45 M using a High Sucrose buffer (2.3 M Sucrose 20 mM HEPES/KOH pH 7.4, 2mM EDTA, 2 mM PMSF). The suspension was sonicated eight times with 10 second pulses, alternating with a break of 1 min on ice. The sonicated sample was clarified (35000 xg, 30 min, 4°C) and the pellet was resuspended in 15 mL of swelling buffer. The solution was centrifuged (200,000 xg, 2 h, 4°C) to harvest the mitochondrial vesicles. The pellet was resuspended in 400 µL of low sucrose buffer (0.45 M Sucrose 20 mM HEPES/KOH pH 7.4, 2 mM EDTA, 2 mM PMSF), pipetted up and down several times, and sonicated in a sonifying bath at 4°C for 5 min.

Next, the sample was clarified (35000 xg, 15 min, 4°C) and the pellet was resuspended in 1.5 mL of swelling buffer. The supernatant (350 µL) was transferred to a new tube and the sucrose concentration was adjusted to 0.85M using the sucrose gradient buffer (2.5 M sucrose, 5 mM MOPS/KOH, 1 mM EDTA, 50 mM KCl). The supernatant was loaded onto a sucrose step gradient assembled using the sucrose gradient buffer and the gradient buffer (5 mM MOPS/KOH, 1 mM EDTA, 50 mM KCl) where each layer has a volume of 625 µL. The gradient was centrifuged in a SW60 rotor (230000 xg, 16 h, 4°C) such that the lighter MOM vesicles float up while the denser MIM vesicles settle at the bottom. After the centrifugation step, fractions (250 µL, each) were collected starting from the top of the gradient, resuspended in sampled buffer, and analysed by SDSPAGE and immunoblotting.

#### Mass spectrometric lipid analysis

Quantitative analysis of lipids was performed by standard nanoelectrospray ionization mass spectrometry (56, 57). Lipids were extracted from 50 µg of yeast mitochondria in the presence of internal standards for the major phospholipid classes and neutral lipids (PC 17:0-14:1, PE 17:0-14:1, PI 17:0-14:1, PS 17:0-14:1, PG 17:0-14:1, 15:0-18:1-d7-PA; Cardiolipin mix I; d5-TG ISTD Mix I; d5-DG ISTD Mix I; d5-DG ISTD Mix II, all from Avanti Polar Lipids). Extraction was performed according to Bligh and Dyer with some optimization for whole yeast cells (52). Lipids were dissolved in 10 mM ammonium acetate in methanol and analyzed in a QTRAP 6500 triple quadrupole mass spectrometer (SCIEX) equipped with a nanoinfusion spray device (TriVersa NanoMate, Advion). Mass spectra were processed by LipidView Software Version 1.2 (SCIEX) for identification and quantification of lipids. Lipid amounts (pmol) were corrected for response differences between internal standards and endogenous lipids.

## Data availability

All data contained within the manuscript.

## Supporting information

This article contains supporting information.

## Author Contributions

VT, TT, and KSD performed experiments and analysed data together with DR. KSD designed the study with advice from DR. VT and KSD wrote the manuscript with advice from TT, MJ, and DR.

## Acknowledgements

We thank E. Kracker for technical assistance and F. Beitel and P. A. Campos for their help during their internships. We would also like to thank T. Langer for providing yeast strains and G. Zocher for helping with the structural analysis.

## Funding and additional information

This work was supported by the Deutsche Forschungsgemeinschaft (DI 1386/2-2 to KSD).

## Conflict of interest

The authors declare that they have no conflict of interest.

## Notes

### Competing Interest Statement

The authors have declared no competing interest.

