## Supporting information for "Characterization of Tgl2, a putative lipase in yeast mitochondria"

Table S1: Strains analysed for changes in the complex and dimer formation of Tgl2

| **Gene** | **Localization** | **Function** |
|---|---|---|
| *MCP2* | MIM | Lipid homeostasis; CoQ mobilization from the MIM |
| *UPS1* | MIM/IMS | Phosphatidic acid transporter involved in cardiolipin metabolism |
| *UPS2* | IMS | Forms a complex with Mdm35 and transports phosphatidylserine from MOM to MIM; involved in phospholipid metabolism |
| *MDM35* | IMS | Forms a complex with Ups2 and transports phosphatidylserine from MOM to MIM |
| *MDM31* | MIM | Potentially involved in phospholipid metabolism |
| *MDM32* | MIM | Potentially involved in phospholipid metabolism |
| *FIS1* | MOM | Involved in mitochondrial fission |
| *FZO1* | MOM | Involved in mitochondrial fusion |
| *UGO1* | MOM | Involved in mitochondrial fusion by facilitating fusion of MIM and MOM |
| *OM45* | MOM | Unknown function |
| *MIR1* | MIM | Transmembrane phosphate transporter |
| *TGL1* | LDs | Steryl ester hydrolase |
| *TGL3* | LDs | TAG lipase and lysophophatidylethnaolamine acyltransferase |
| *GEP4* | Matrix | Phosphatidylglycerophosphatase, involved in Cardiolipin metabolism |
| *CRD1* | MIM | Cardiolipin synthase, involved in Cardiolipin metabolism |
| *TAZ1* | MOM | Lysolecithin acyltransferase, involved in phospholipid metabolism and cardiolipin remodeling |
| *PSD1* | MIM/ER | Phosphatidylserine decarboxylase, involved in phospholipid metabolism and LD formation |
| $RHO^0$ | - | Yeast strain depleted of mtDNA |

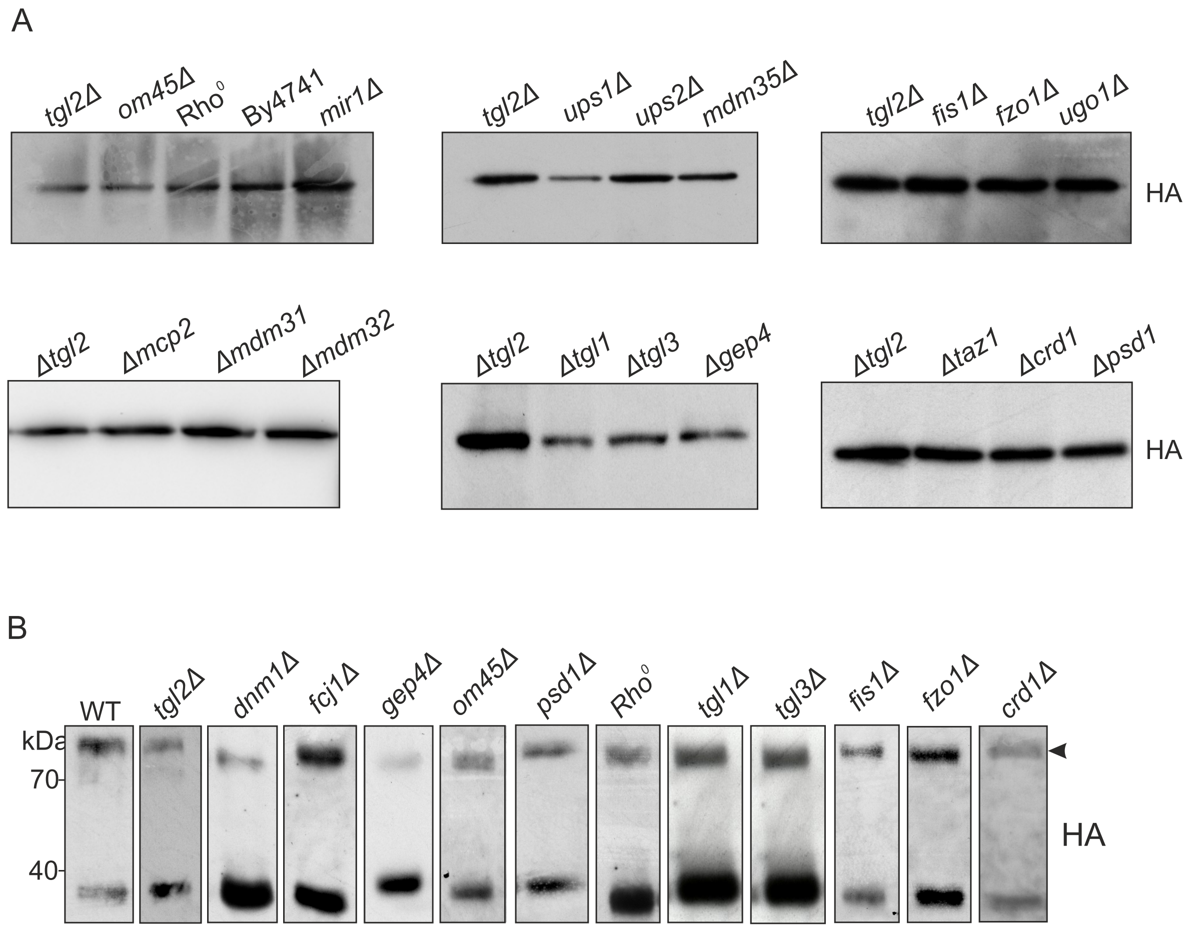


**Fig. S1 Tgl2 complex in different deletion mutants**
(A) The Tgl2 complex is unaffected in the absence of the indicated strains. Mitochondria were isolated from the mentioned strains, solubilized with digitonin, and analysed by BN-PAGE (4-14%) and immunodecoration against HA-tag. .There were noticeable effects on the complex formation. (B) Steady state levels of Tgl2 in different deletion strains. Mitochondria were isolated from the indicated strains expressing HA-Tgl2 and analysed by SDS-PAGE and immunodecoration with an antibody against HA-tag. The Western blots are representative of some strains used in this experiment. The intensity of the dimer was quantified with respect to total amount of Tgl2 (n = 3).

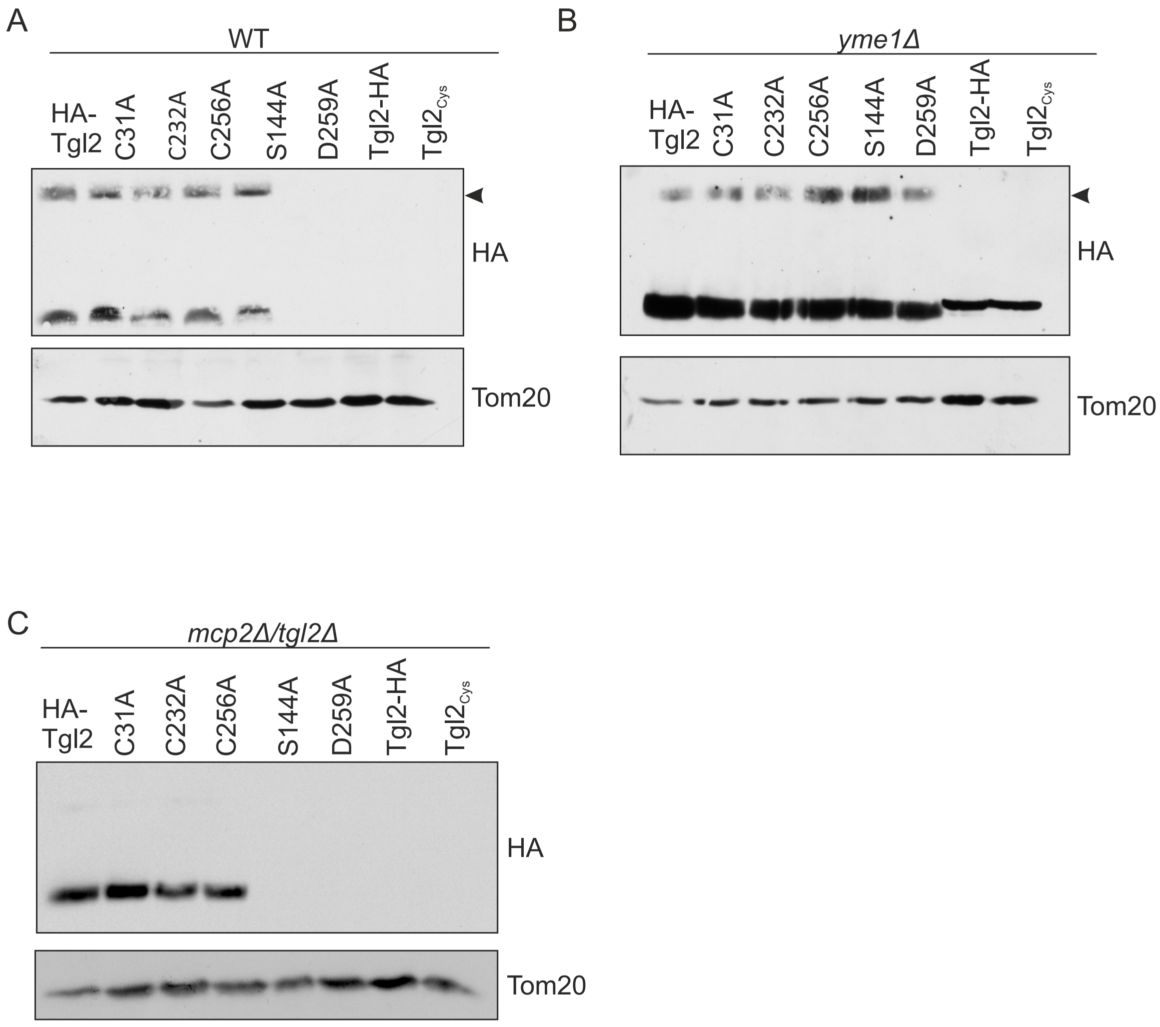


**Fig. S2 Steady state levels of Tgl2 mutants**

(A-C) The expression Tgl2 mutants varies in WT, *yme1*, and *mcp2/tgl2* cells. Cells expressing the the indicated variants of Tgl2 were grown to mid-logarithmic phase and crude mitochondrial fractions were isolated. The organelles were analysed by SDS-PAGE and immunodecoration with antibodies against the HA-tag and Tom20, as a loading control. All variants of Tgl2 can be detected in *yme1* cells. Tgl2HA, Tgl2$_{cys}$, and HA-Tgl2$_{S144A}$ can only be detected in WT cells. HA-Tgl2$_{D259A}$is detectable in WT cells only in higher exposures (Fig. 7C). In *mcp2 /tgl2,* only the functional variants can be detected at comparable levels to the native protein. The non-functional variants are detectable at longer exposures.

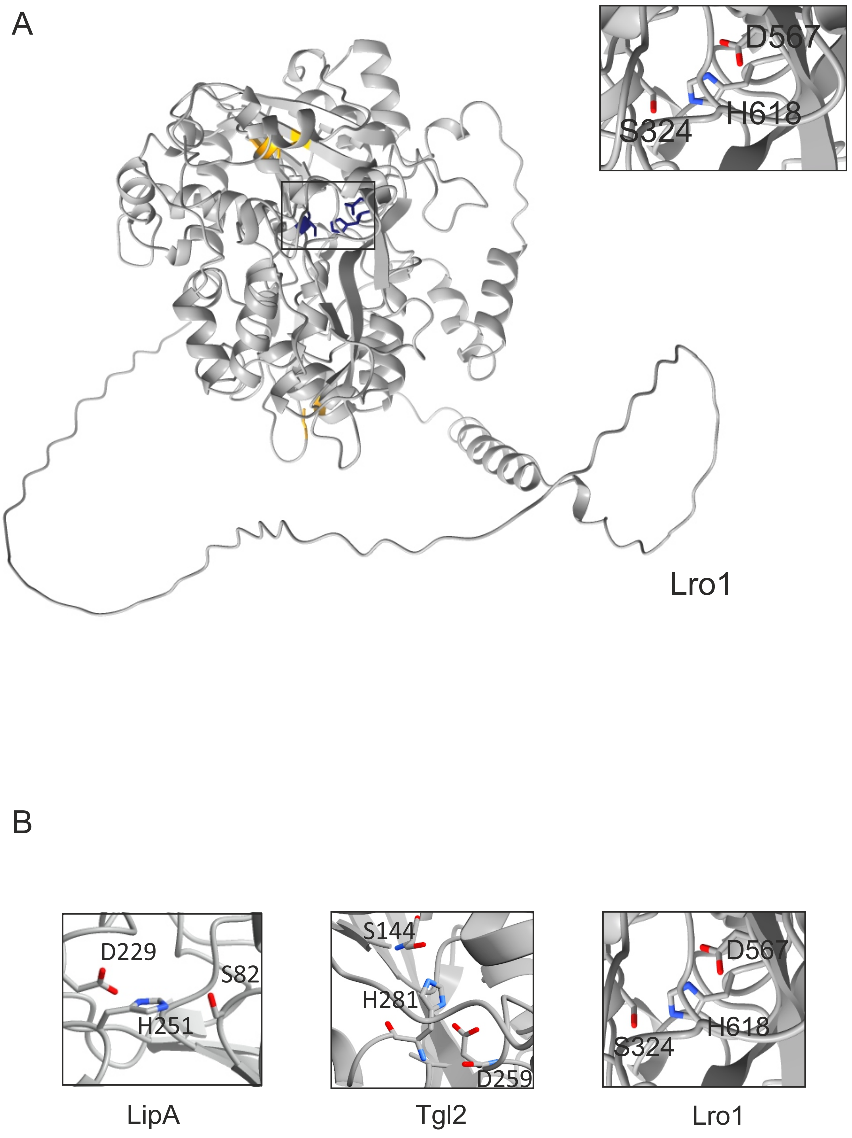


**Fig. S3 Catalytic triad of Lro1 – a yeast acyltransferase**
(A) Acyltransferases contain a catalytic triad consisting of Ser-Asp-His. Predicted structure of Lro1 with the intramolecular disulfide bonds in yellow and the catalytic triad highlighted in blue. (B) The Ser-Asp-His catalytic triad of LipA (left, a bacterial lipase), Tgl2 (center, yeast putative TAG lipase), and Lro1 (right, yeast acyltransferase).
